# Interrogating the genus *Yersinia* to define the rules of lipopolysaccharide lipid A structure associated with pathogenicity

**DOI:** 10.1101/2025.03.03.641127

**Authors:** Maria Helena Fernandez-Carrasco, Käthe M. Dahlström, Ricardo Calderon-Gonzalez, Amy Dumigan, Jan K. Marzinek, Rebecca Lancaster, Peter J. Bond, Tiina A. Salminen, José A. Bengoechea

**Author notes:** Corresponding author (J.A. Bengoechea).

## Abstract

Pathogen recognition by the immune system relies on germline-encoded pathogen recognition receptors which identify conserved pathogen-associated molecular patterns (PAMPs) such as the lipid A section of the lipopolysaccharide (LPS). The assumption that pathogens and mammalian-associated bacteria remodel their lipid A PAMP because of host-microbe co-evolution is a long held-belief of microbial pathogenesis. We set out to test this fundamental principle by interrogating a Gram-negative genus presenting evidence of evolutionary events linked to the acquisition of essential virulence traits, resulting in pathogenic and non-pathogenic species. The genus *Yersinia* fulfil these requirements; the acquisition of the pYV virulence plasmid is one of the evolutionary events associated with virulence. At 37°C, only pathogenic *Yersinia* switch to deacylated lipid A, a modification that diminishes TLR4/MD-2 recognition and reduces inflammation. An engineered chimeric pathogenic *Yersinia* strain expressing the non-pathogenic lipid A profile efficiently engages with TLR4, demonstrating it is sufficient to switch the acylation pattern to modify the recognition by TLR4 and subsequent activation of inflammation. The lipid As of pathogenic and non-pathogenic species are modified with aminoarabinose and palmitate; therefore, only the reduced acylation of the lipid A PAMP is a trait associated with virulence. The decorations of lipid A do not alter TLR4 engagement but confer resistance to antimicrobial peptides. The chimeric pathogenic *Yersinia* strain expressing the non-pathogenic lipid A profile allows to ascertain whether the switch in the lipid A PAMP affects virulence. This strain showed enhanced motility due to an upregulation of the *flhDC* master regulator, and impaired cellular invasion through downregulation of *rovA*, a key invasion regulator. The expression and function of pYV-encoded virulence factors Yops and YadA were not affected. Nonetheless, the chimeric strain was attenuated in vivo, demonstrating that virulence factors cannot overcome a switch in the lipid A PAMP associated with pathogenicity.

## INTRODUCTION

The recognition of pathogens through the actions of a set of germline encoded innate immune receptors is one of the foundations of immunology. These receptors, known as pattern recognition receptors (PRRs), detect conserved molecules common to large number of microbes to launch an antimicrobial programme [1]. These microbial molecules are distinguishable from “self”, important for the viability and fitness of the pathogen in different environments and, therefore, they are stable [2]. These molecules contain so-called pathogen-associated molecular patterns (PAMPs) recognized by the PRRs [2]. Well-characterized molecules are the lipid A section of the lipopolysaccharide (LPS), the flagellin of the bacterial flagella, and bacterial nucleic acids.

The limits of the universal rule of PRR recognition of PAMPs were challenged when it was recognized that pathogens and members of the microbiome may modify their PAMPs to affect the activation of PRRs [3–5]. This has been particularly studied in the case of the LPS lipid A recognized through the PRRs Toll-like receptor 4/myeloid differentiation factor 2 (TLR4/MD-2) complex, and inflammatory caspases, caspase-11 in mouse and caspases-4/5 in humans [6–10]. LPS receptors best sense a bisphosphorylated diglucosamine to which are attached six saturated fatty acyl chains with lengths of 12 or 14 (occasionally 16) carbons, the so-called “hexa-acyl lipid As” [11]. Variations of this structure by changing the number and length of fatty acids, the phosphorylation status as well as by adding some lipid A decorations, phosphoethanolamine and aminoarabinose may result in impaired PRR recognition [12]. Therefore, these changes of the lipid A PAMP are thought to represent adaptations of microbes to the challenge imposed by the innate immune system as a result of the co-evolution between host and microbes. However sound this notion, much of the research has focused only on pathogens, and there is extensive evidence showing that environmental conditions influence the lipid A structure. It cannot then be rigorously ruled out that these lipid A changes found in pathogens may simply reflect bacterial adaptation to changing environments and, therefore, are shared by non-pathogenic Gram-negative bacteria. Supporting this notion, deep-sea bacteria express lipid A with decorations found in pathogenic bacteria [13]. Therefore, the question remains whether indeed there is a distinct lipid A PAMP expressed by pathogens shaped by host-pathogen co-evolution.

To test this principle rigorously, it is necessary to interrogate bacteria from the same genus as they should produce the same lipid A. Within the genus, there should be evidence of evolutionary events linked to the acquisition of essential virulence traits, resulting in, at least, two well defined distinct groups, one encompassing pathogenic species and another including non-pathogenic species lacking almost all the virulence factors expressed by the pathogenic group. The comparison between these two groups of species should help to delineate features of the lipid A PAMP of pathogens if they do exist.

Careful consideration of Gram-negative bacterial diversity reveals that the genus *Yersinia* meets all the requirements to define the lipid A PAMP associated with virulence. The genus includes 26 species of which only *Y. pestis*, and the enteropathogenic *Y. pseudotuberculosis* and *Y. enterocolitica* are mammalian (including humans) pathogens [14, 15]. The remaining known species are commonly found in soil and aquatic environments and are non-pathogenic in mammals [16]. Evolution studies of the genus *Yersinia r*evealed that the most prominent event during the emergence of pathogenesis is the acquisition of the pYV virulence plasmid, encoding the Ysc type III secretion system (T3SS) crucial to blunt cell-intrinsic immunity [14, 15, 17].

In this study, by examining pathogenic and non-pathogenic species of the genus *Yersinia*, we demonstrate that indeed there is a lipid A PAMP associated with virulence. This lipid A PAMP is characterized by reduced acylation. In contrast, the decorations of the lipid A PAMP with aminoarabinose and palmitate, though useful to survive within the mammalian host, are not unique traits of the pathogenic lipid A PAMP. We provide molecular mechanistic insights into the enzymology of the lipid A PAMP. Finally, our results demonstrate the negative effect on virulence of switching the lipid A PAMP expressed by pathogens.

## RESULTS

### Variations in the lipid A PAMP expressed by pathogenic and non-pathogenic *Yersinia spp*

In *Y. pestis*, and the highly virulent enteropathogenic *Y. pseudotuberculosis* and *Y. enterocolitica* phylogoup 2 (serotype O:8) (hereafter YeO8), growth temperature regulates the lipid A acylation pattern being predominantly hexa-acylated and hepta-acylated at 21°C and tetra-acylated at 37°C [18–22]. Only in bacteria grown at 21°C, the lipid A is decorated with aminoarabinose and palmitate [18, 20, 23–25]. To investigate whether this lipid A PAMP is found in other pathogenic *Yersiniae*, we determined the lipid A structure of two additional enteroptahogenic *Y. enterocolitica* strains from phylogroup 3 (serotype O3, hereafter YeO3) and phylogroup 5 (serotype O:9, hereafter YeO9) (Table S2). Phylogroup 3 is the most frequent cause of human yersiniosis [26]. We extracted lipid A and predicted structural composition by MALDI-TOF mass spectrometry (MS) in the negative ion mode. Mass spectral data were used to predict the number of acyl changes, the loss of a phosphate from the di-glucosamine sugar backbone (mono-phosphorylation), and the decorations of the lipid A with aminoarabinose, palmitate, or phosphoethanolamine. When grown at 21°C, YeO3 and YeO9 lipid As contained primary ionizable peaks *m/z* 1,797 and 1,824 (Fig 1A and Fig 1B), which are predicted to be hexa-acylated lipid A species (Table 1). *m/z* 1,824 corresponds to a structure containing two glucosamine residues, two phosphate groups, four 3-OH-C14, one C12 (laurate), and one C16:1 (palmitoleate) (Table 1). *m/z* 1,797 contains C14 (myristate) instead of C16:1. Peak *m/z* 1,414 corresponds to a tetra-acylated lipid A (Table 1); we have demonstrated that this species is caused by the activity of the LpxR deacylase that removes the 3′-acyloxyacyl residue of the lipid A [20]. YO3 and YeO9 also encode the *lpxR* deacylase (Table S1). Two more peaks of *m/z* 1,954 and 2,063 were detected and they correspond to the modification of the species *m/z*1824 with aminoarabinose (*m/z* 131) and palmitate (*m/z* 239), respectively (Fig 1A and Fig 1B). YeO3 and YeO9 encode the *pmr/arn* operon responsible for the decoration with aminoarabinose, and the *pagP* acyltransferase responsible for the addition of palmitate to produce hepta-acylated lipid A (Table S1) [23]. These lipid A species are also found in the lipid A of YeO8 grown at 21°C (Table 2) [20, 21, 23]. When grown at 37°C, YeO3 and YeO9 lipid As contained hexa-acylated lipid A, *m/z* 1,797, and a tetra-acylated lipid A, *m/z*1,388, corresponding to a species with two glucosamine residues, two phosphate groups, three 3-OH-C14 units, and one C14 (Fig 1C and Fig 1D). These lipid A species are also found in the lipid A of YeO8 grown at 37°C [20] (Table 2). *m/z* 1,388 peak is consequence of the action of LpxR [20]. No lipid A modifications were detected (Fig 1C and Fig 1D). Altogether, it can be concluded that the lipid A PAMP of pathogenic *Yersinia spp* is characterized by its reduced acylation at 37°C and by the temperature-dependent decoration of lipid A with aminoarabinose and palmitate.

**Figure 1.**
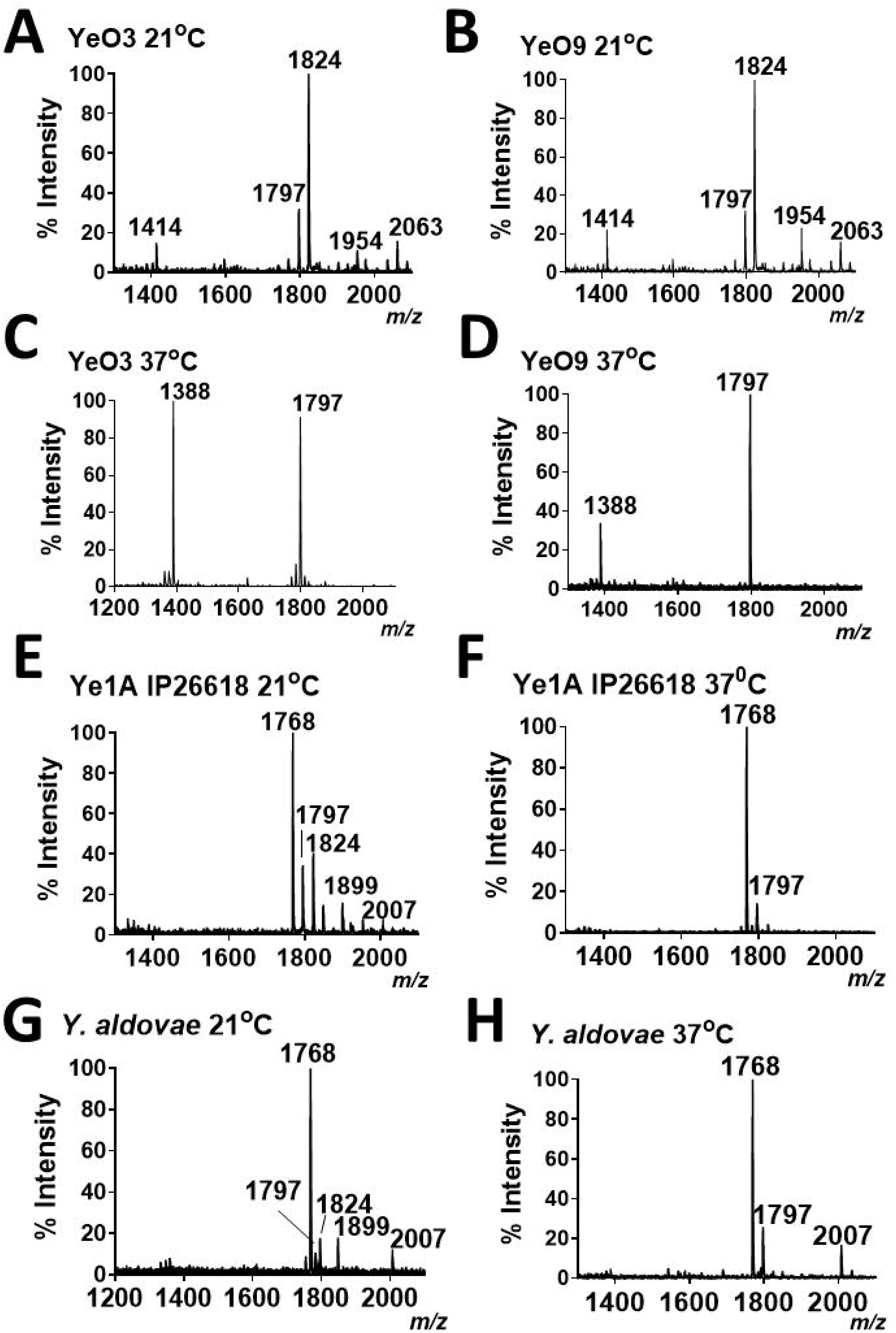
Characterization of *Yersiniae* lipid A. Negative ion MALDI□TOF mass spectrometry spectra of lipid A purified from (A) YeO3 grown at 21°C, (B) YeO9 grown at 21°C, (C) YeO3 grown at 37°C, (D) YeO9 grown at 37°C, (E) Ye1A strain IP266118 grown at 21°C, (F) Ye1A strain IP266118 grown at 37°C, (G) *Y. aldovae* grown at 21°C, and (H) *Y. aldovae* grown at 37°C.. Data represent the mass□to□charge (*m/z*) ratios of each lipid A species detected and are representative of three extractions.

To ascertain the lipid A PAMP expressed by non-pathogenic *yersiniae*, we analysed the lipid A from five strains belonging to *Y. enterocolitica* phylogroup 1A (Table S2). This phylogroup includes non-pathogenic *Y. enteroclitica* strains isolated from a variety of environmental sources and they do not harbour the pYV virulence plasmid [17, 27]. Strikingly, these strains expressed a different lipid A than the pathogenic *Y. enterocolitica* strains (Table 2). When grown at 21°C, we detected peaks *m/z* 1,797 and 1,824 (Fig 1E and Table 1) and a new species *m/z* 1,768. The latter corresponds to a hexa-acylated lipid A containing two glucosamine residues, two phosphate groups, four 3-OH-C14, and two C12 (laurate) (Table 1). This lipid A species was modified with aminoarabinose, *m/z* 1,899, and palmitate, *m/z* 2,007 (Fig 1E and Table 1). When grown at 37°C, these non-pathogenic strains expressed hexa-acylated lipid As *m/z* 1768 and *m/z* 1,797, and no decorations were detected (Fig 1F and Table 2). No tetra-acyl lipid A species were detected at any growth temperature. These results suggest that the reduced acylation of the lipid A at 37°C could be a trait of the lipid A PAMP from pathogenic strains whereas this is not the case for the decorations of the lipid A with aminoarabinose and palmitate.

We next extracted lipid As from twelve non-pathogenic *Yersinia spp*, *Y. aldovae, Y. bercovieri, Y. mollaretii, Y. frederiksenii, Y.intermedia, Y. kristensenii, Y. rhodei, Y. ruckeri, Y. similis, Y. massilensis, Y. pekkaneniii,* and *Y. nurmii* (Table S2). The lipid As extracted from these species grown at 21°C were similar to those produced by *Y. enterocolitica* 1A strains (Fig 1G and Table 2). At 37°C, we did not detect any tetra-acylated lipid A species (Fig 1H and Table 2), and the peaks detected corresponded to hexa-acylated lipid As *m/z* 1,768 and 1,797 (Table 1). Only for *Y. aldovae, Y. bercovieri, Y. mollaretii, Y. frederiksenii, Y. intermedia,* and *Y. kristensenii* lipid As did we detect a peak *m/z* 2,007, consistent with the addition of palmitate to the hexa-acylated species *m/z* 1,768 (Fig 1H, Table 1 and Table 2).

Altogether, these results demonstrate that the switch to deacylated lipid As at 37°C only occurs in pathogenic *Yersiniae*. The temperature-dependent variation in the modifications of the lipid A observed in pathogenic *Yersiniae* is also found in non-pathogenic *Yersiniae* (this work) [18, 20, 23–25]. Therefore, the modifications of the lipid A with aminoarabinose and palmitate cannot be considered a trait uniquely found in the lipid A PAMP of pathogenic bacteria in contrast to the reduced acylation.

### Enzymology of *Yersinia spp* lipid A PAMP acylation

One conspicuous difference between the lipid As of the pathogenic and non-pathogenic *Yersiniae* is the absence of tetra-acylated species in the latter. Bioinformatic analysis revealed the absence of the deacyase *lpxR* in the genomes of the non-pathogenic *Yersiniae* (Table S1), explaining the lack of LpxR-dependent tetra-acylated lipid As *m/z* 1,388 and 1,414 in the lipid As of these species. We questioned whether LpxR will be sufficient to produce deacylated lipid As in non-pathogenic *Yersiniae*. YeO8 *lpxR* was introduced at a single copy in *Y. aldovae* genome using the Tn7 delivery system [28]. When grown at 21°C, the lipid A expressed by this chimeric non-pathogenic strain, *Y. aldovae*::LpxR_YeO8_, contained peaks *m/z* 1,414 and 1,388 consistent with the deacylation of the hexa-acylated species *m/z* 1,797 and *m/z* 1,824 (Fig 2A). At 37°C, we detected lipid A species *m/z* 1,360 and *m/z* 1,388 resulting from the de acylation of the hexa-acylated species *m/z* 1.768 and 1,797, respectively (Fig 2B). These results demonstrate that the lack of lipid A deacylation is not an intrinsic property of non-pathogenic *Yersiniae* lipid As.

**Figure 2.**
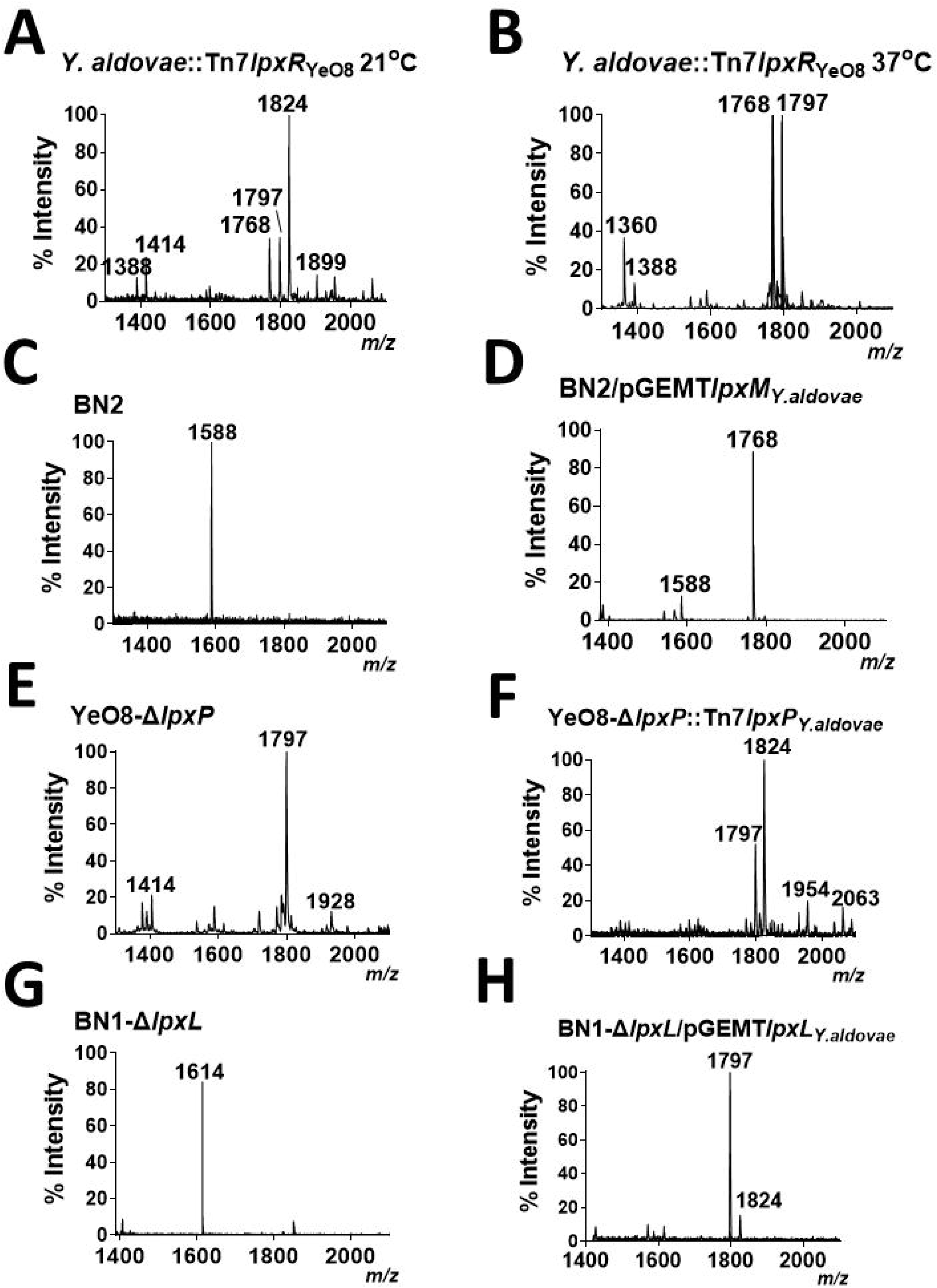
Structural characterization of *Yersiniae* lipid A enzymes. Negative ion MALDI□TOF mass spectrometry spectra of lipid A purified from (A) *Y. aldovae* expressing *lpxR* from YeO8 grown at 21°C, (B) *Y. aldovae* expressing *lpxR* from YeO8 grown at 37°C, (C) YeO3 grown at 37°C, (C) *E. coli* BN2, (D) *E. coli* BN2 complemented with *lpxM* from *Y. aldovae*, (E) YeO8 *lpxP* mutant grown at 21°C, (F) YeO8 *lpxP* mutant complemented with *lpxP* from *Y. aldovae* grown at 21°C, (G) *E. coli* BN1 *lpxL* mutant, (H) *E. coli* BN1 *lpxL* mutant complemented with *lpxL* from *Y. aldovae*. Data represent the mass□to□charge (*m/z*) ratios of each lipid A species detected and are representative of three extractions.

Another conspicuous difference is the presence of the hexa-acylated species *m/z* 1,768 only in the lipid As from non-pathogenic *yersiniae* (Table 2). In pathogenic *Yersiniae,* the late acyltransferase LpxM transfers C12 to the 3′-R-3-hydroxymyristoyl group, whereas LpxL and LpxP transfer C14 and C16:1, respectively to the 2′-R-3-hydroxymyristoyl group [21, 22]. In silico analysis revealed the genomes of non-pathogenic *Yersiniae* encoded three late acyltransferases, *lpxM, lpxL* and *lpxP* (Table S1) with more than 90% sequence identity to the pathogenic late-acyltransferases. We then sought to characterize the substrate specificity of these enzymes in vivo by expressing them in a surrogate host. This approach has been used previously to characterize the function of lipid A enzymes in the context of the bacterial membrane [29, 30]. *Y. aldovae lpxM* was cloned and expressed in *E. coli lpxM* mutant strain BN2 [31]. As expected, lipid A extracted from BN2 contained only one species *m/z* 1,588 corresponding to two glucosamine residues, two phosphate groups, four 3-OH-C14, and one C12 (added by the *E. coli* acyltransferase LpxL) (Fig 2C). BN2 complemented with *Y. aldovae lpxM* contained species *m/z* 1,768 consistent with the addition of C12 to the lipid A (Fig 2D), establishing that LpxM from non-pathogenic *Yersiniae* esterifies the lipid A with C12 likewise LpxM from pathogenic *Yersiniae* [21, 22]. Similar results were obtained expressing *lpxM* from *Y. bercovieri*, and *Y. intermedia (*Fig S1A and Fig S1B).

To confirm whether LpxP from non-pathogenic *Yersiniae* transfers C16:1 to 2′-R-3- hydroxymyristoyl group like LpxP from pathogenic *Yersiniae* [21, 22], we complemented YeO8 *lpxP* mutant, strain YeO8-Δ*lpxPGB* [21], with *lpxP* from *Y. aldovae. lpxP* from the non-pathogenic strain restored the species *m/z* 1,824 in YeO8 *lpxP* mutant (Fig 2E and Fig 2F); demonstrating that LpxP from non-pathogenic *Yersiniae* transfer C16:1 to the lipid A. *Y. bercovieri* and *Y. intermedia lpxP* also complemented YeO8-Δ*lpxPGB* (Fig S1C and Fig S1D).

These enzymatic activities raise the question which is the late acyltransferase esterifying the 2′-R-3- hydroxymyristoyl group with C12 to give *m/z* 1,768 in non-pathogenic *Yersiniae*. It can be postulated that either LpxL from non-pathogenic *Yersiniae* has a relaxed specificity and can transfer C12 and C14 to the 2′-R-3-hydroxymyristoyl group, or non-pathogenic *Yersiniae* encode a hitherto unknown late acyltransferase. Ruling out the latter, analysis of the genomes of non-pathogenic *Yersiniae* did not reveal the presence of another putative late lipid A acyltransferase. To determine whether LpxL from non-pathogenic *Yersiniae* has a relaxed specificity, we determined the activity of LpxL in the background of *E. coli lpxL* mutant, strain BN1-Δ*lpxL* [30]. This strain produces a penta-acylated lipid A species, *m/z* 1,641 (Fig 2G) containing two glucosamine residues, two phosphate groups, four 3-OH-C14, and one C14 (added by *E. coli* LpxM) [30]. The lipid A produced by BN1-Δ*lpxL* harbouring LpxL from YeO8 contained species *m/z* 1,824, consistent with the addition of C14, confirming previous findings (Fig S1E). [21]. Notably, BN1-Δ*lpxL* expressing LpxL from *Y. aldovae* produced lipid As containing species *m/z* 1,797, consistent with the addition of C12, and *m/z* 1,824, consistent with the addition of C14 (Fig 2H). Similar results were observed when *lpxL* from *Y. bercovieri,* and *Y. mollaretii* were expressed in BN1Δ*lpxL* (Fig S1F and Fig S1G). Altogether, these results establish that LpxL from non-pathogenic *yersiniae* esterifies the lipid A 2′-R-3-hydroxymyristoyl group with C12 and C14, demonstrating a relaxed acyl chain length selectivity of lipid A late acyltransferases.

### Molecular basis of non-pathogenic *Yersinia spp* LpxL relaxed specificity

To elucidate the molecular basis of the relaxed specificity for C12 and C14 observed in non-pathogenic LpxL, we used template-based homology modeling to predict the structural models for LpxL from YeO8, *Y. aldovae*, and *E. coli.* The models were based on the crystal structure of *Acinetobacter baumannii* LpxM (PDB ID 5KN7) [32], which showed up as a relevant hit (E-value of 3e-12) when searching PDB with the YeO8 LpxL sequence as bait. LpxM, LpxL and glycerol-3- phosphate acyltransferases (GPAT) all belong to the lysophospholipid acyltransferases (LPLATs) superfamily, and share the GPAT family motifs (cyan in Fig S2). The crystal structure was used to identify the active site (corresponding to the LpxM site with bound n-dodecyl-β-D-maltoside) with an entrance to a deep hydrophobic tunnel (corresponding to the LpxM tunnel with a glycerol bound) in the studied LpxL proteins. Due to its hydrophobic nature, this tunnel was predicted to serve as the binding site for an acyl chain (Fig 3). At the bottom of the tunnel, YeO8 LpxL has a leucine (L146), which corresponds to a phenylalanine in *Y. aldovae* (F142) and *E. coli* (F143) LpxL. This residue likely functions as a hydrocarbon ruler and confers the ability to precisely measure and incorporate hydrocarbon chains of a specific length [33, 34], similarly to residues in enzymes from the GPAT family. Homology models of *E. coli*, *Y. aldovae* and YeO8 LpxL with C12 and C14 docked to the hydrophobic tunnel predict a stable complex between *E. coli* LpxL and C12, whereas the binding of C14 is not hampered by the side chain of F143 restricting the tunnel size (Fig 3). Moreover, F143 is unlikely to swing away to make space for C14 due to the more polar environment at the bottom of the tunnel, which is unfavourable for the hydrophobic side chain of F143 (Fig S3). Also *Y. aldovae* LpxL is predicted to bind C12, with F142 stabilizing the interaction with the acyl chain. However, *Y. aldovae* LpxL F142 may be able to swing away from the tunnel cavity to provide a stable interaction site also with C14, since the surrounding amino acids L112, I141 and L253 are small hydrophobic residues, which can interact with both F142 and the carbon tail of C14. On the other hand, YeO8 LpxL L146 has a smaller side chain than phenylalanine, thereby facilitating and stabilizing the binding of C14, but being unable to form interactions with the shorter C12 that would place the acyl chain in the correct position for the catalysis to take place. Altogether, these findings suggest that F143 (*E. coli* LpxL), F142 (*Y. aldovae* LpxL) and L146 (YeO8 LpxL) act as hydrocarbon rulers. Furthermore, the alignment of *Yersinia* LpxL homologs showed a high conservation rate among the residues lining the hydrophobic tunnel, while also highlighting that the hydrocarbon ruler residue is conservatively a phenylalanine in non-pathogenic species and a leucine in pathogenic species (Fig S2).

**Figure 3.**
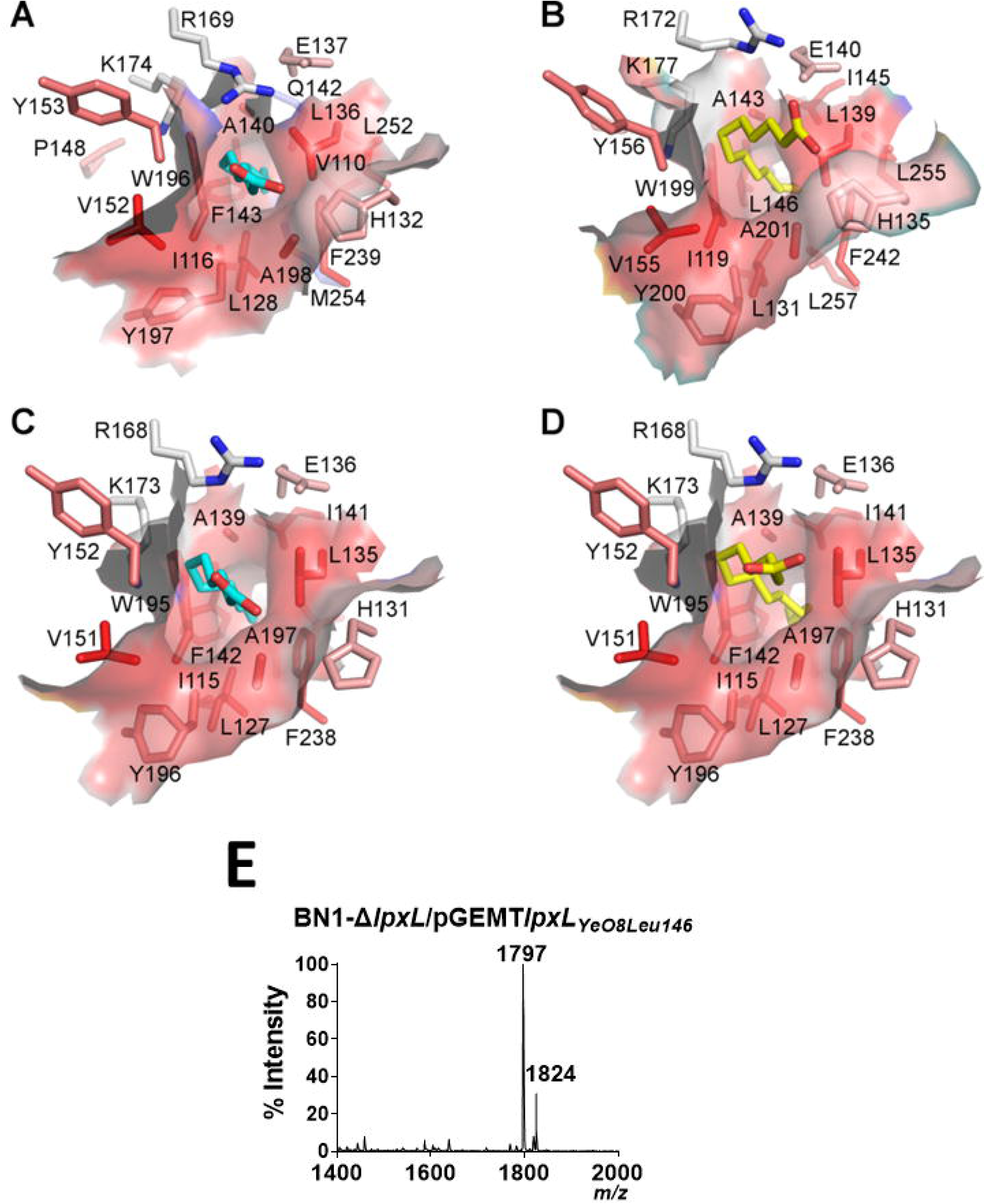
Defining *Yersiniae* LpxL hydrophobic tunel. A deep hydrophobic tunnel is formed in all LpxL models and may serve as the binding site for an acyl chain. The surface of the tunnel is color coded by hydrophobicity in PyMOL, where gray shows charged residues and red hydrophobic residues. In all models, the amino acids contributing to the tunnel are mostly hydrophobic. (A) *E. coli* LpxL with C12 (cyan sticks). (B) YeO8 LpxL with C14 (yellow sticks), (C) *Y. aldovae* LpxL with C12 (cyan sticks), (D) *Y. aldovae* LpxL with C14 (yellow sticks). (E) Negative ion MALDI□TOF mass spectrometry spectra of lipid A purified from *E. coli* BN1 *lpxL* mutant complemented with *lpxL* from YeO8 in which L146 is mutated to phenylalanine. Data represent the mass□to□charge (*m/z*) ratios of each lipid A species detected and are representative of three extractions.

To test these predictions, we mutated the putative hydrocarbon rule of YeO8 LpxL from leucine to phenylalanine and assessed the specificity of the enzyme in vivo. When expressed in BN1-Δ*lpxL* strain, the lipid A contained species *m/z* 1,797 and *m/z* 1,824 (Fig 3E), validating that L146 is the hydrocarbon ruler of YeO8 LpxL and that its replacement by phenylalanine is sufficient to relax the acyl chain specificity of the enzyme.

### Molecular dynamics simulations of TLR4_2_/MD-2_2_ with the lipid A PAMP

We next investigated the ability of mammalian LPS receptors to interact with the lipid A PAMP produced by pathogenic and non-pathogenic *Yersiniae*. Although TLR4 is often considered the receptor for LPS, MD-2, a co-receptor that physically interacts with the extracellular domain of TLR4, is principally responsible for LPS binding [35]. In the inactive state, TLR4 exists as a heterodimer with the co-receptor MD-2. The binding of LPS to TLR4/MD-2 creates a dimerization interface that facilitates the formation of the active complex consisting of a “dimer of dimers” (TLR4_2_/MD-2_2_) [35]. LPS interacts with a large hydrophobic pocket in MD-2 and directly bridges the two components of the multimer. The F126 loop of MD-2 is particularly important in stabilizing the active TLR4_2_/MD-2_2_ complex, undergoing a structural shift upon LPS binding [35].

To gain insights into the potential of the different lipid A molecular species as TLR4/MD-2 agonists, we used explicitly solvated atomic-resolution molecular dynamics (MD) simulations to assess the interaction between the lipid A molecular species and the TLR4_2_/MD-2_2_ complex, and with the F126 residues within the MD-2 loop encompassing 120 to 129 residues. We performed six sets of independent 1,000 ns simulations, modelled using the X-ray structure of the hexa-acylated LPS-bound-(TLR4/MD-2)_2_ dimer in its “activated state” (Fig 4A). Each system thus corresponded to the dimeric TLR4/MD-2 complex with two lipid A species, one embedded within each of MD-2 monomers. As a control, we tested *E. coli* lipid A, known to strongly activate the TLR4/MD-2 complex and, therefore, a benchmark molecule to determine activation. Of the *Yersinia* lipid A species, we assessed the hexa-acylated lipid A species *m/z* 1,797 and *m/z* 1,824 found in pathogenic and non-pathogenic *Yersiniae,* the hexa-acylated species *m/z* 1,768 found only in non-pathogenic *Yerisiniae*, the hepta-acuylated lipid A species *m/z* 2,007 found in pathogenic and non-pathogenic *Yersiniae*, and the tetra-acylated lipid A species *m/z* 1,388 present only in pathogenic *Yersiniae* at 37°C (Table 1 and Table 2). We reasoned that these lipid As would allow us to determine the effect of lipid A acylation on the interaction with the TLR4/MD-2 complex and to investigate the effect of the decoration with palmitate. Calculation of the root-mean-square deviation (RMSD) of the TLR4 ectodomains relative to their starting structures over the last 200 ns of simulations revealed that all systems studied, apart from the lipid A species *m/z* 1388, retained the near-experimental “activated state” structure (Fig 4B). Lipid A *m/z* 1.388 amongst all studied species has only four lipid tails in comparison to six or seven held by other species. Hence, the buried area between the MD-2:lipid A complex and the TLR4 dimer as well as total number of contacts between lipid A and (TLR4/MD- 2)_2_ were the lowest in the case of *m/z* 1388 (Fig 4C and Fig 4D). We next looked at the loop of MD- 2 containing F126 which is essential in maintaining the active TLR4_2_/MD-2_2_ complex and overlaid the F126 side chain every 200 ns using two aligned trajectories composed of MD-2:lipid A. In systems containing *E. coli* lipid A or *m/z* 2,007, the F126 side chain retained its orientation facing the hydrophobic lipid tails throughout the entire trajectory for both monomers (Fig 4E). In contrast, lipid A species *m/z* 1,768, *m/z* 1,797 and *m/z* 1,824 retained the F126 side chain orientation in only one of the MD-2 monomers, while *m/z* 1388 did not show a structurally well-defined side chain conformation (Fig 4E). The loop 120-129 containing this residue was also the least stable in the case of *m/z* 1388, whereas no differences were found between any of other lipid A species and *E. coli* (Fig 4F). Altogether, these simulations uncover a reduced interaction of lipid A species *m/z* 1,388 with the TLR4/MD-2 complex, resulting in loss of receptor stability.

**Figure 4.**
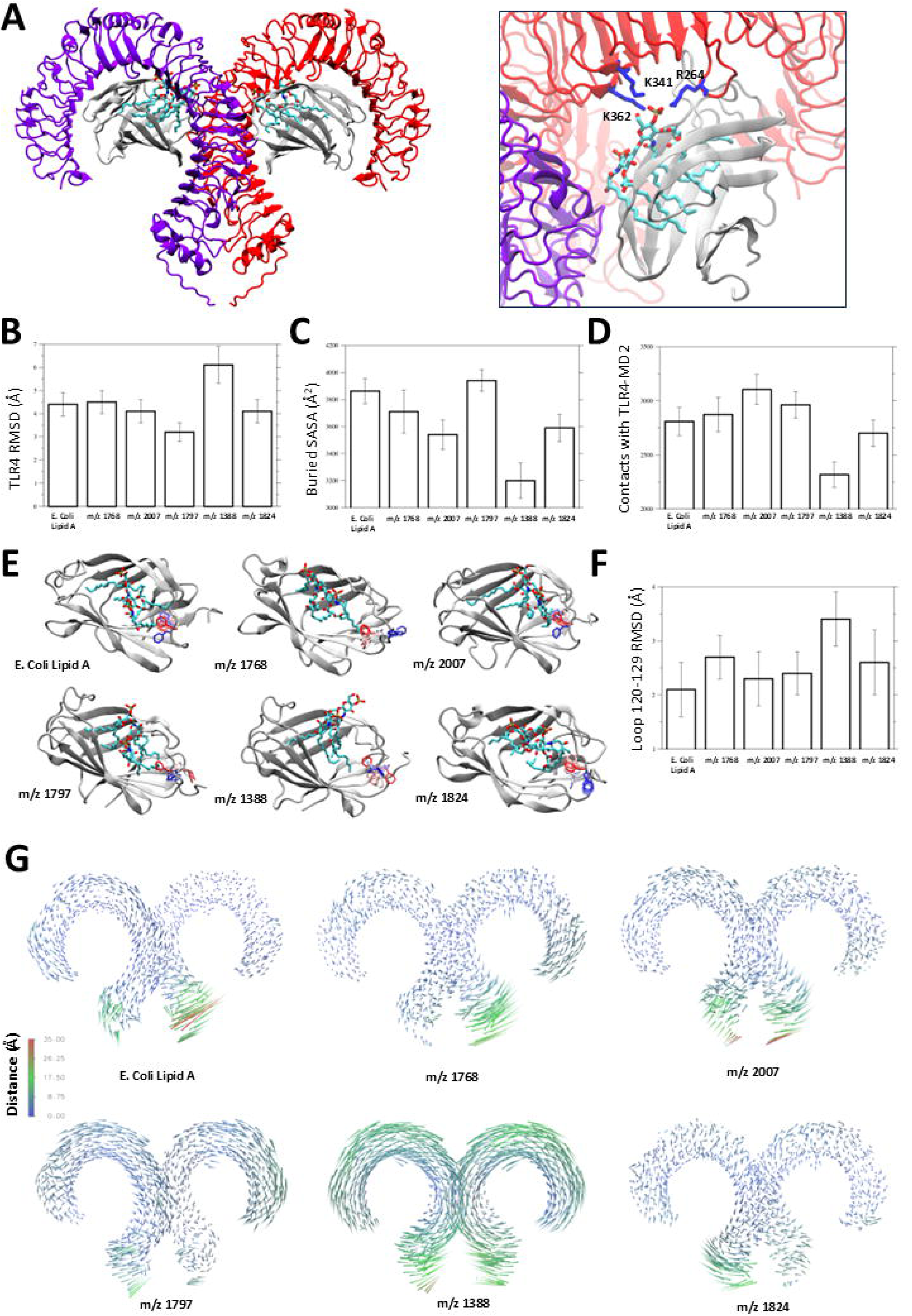
Molecular dynamics simulations of TLR4_2_/MD-2_2_ heterodimer with lipid A species. (A) Initial snapshot of the (TLR4/MD-2)_2_ complex bound to *m/z* 1768 (left). Final snapshots of each one side of the (TLR4/MD-2)_2_ complex bound to *m/z* 1768 (right). Labelled basic amino acid residues observed to make significant hydrogen-bonding interactions with *m/z* 1768 are shown in licorice format, coloured in blue. (B) Root mean square deviation (RMSD) of TLR4 dimer backbone atoms. (C) Buried solvent accessible surface area (SASA) between MD-2-lipid A species and TLR4. (D) Contacts between lipid A species and (TLR4/MD-2)_2_. (E) Overlaid every 200 ns simulation snapshots of F126 side chain and its orientation with respect to the MD-2:lipid A species. The pseudo trajectory was made of two aligned MD-2:lipid monomer frames for each system. The F126 side chain is shown in licorice representation and coloured according to simulation time (red-white-blue for 0 to 2,000 ns). (F) RMSD of MD-2 backbone loop region after alignment on the entire MD-2 backbone. (G) Porcupine plot of TLR4 dimer alpha carbons corresponding to the most dominant motion (PCA1). The colours corresponds to distances as labelled inset. In panel (A) and (E) protein is shown in cartoon representation: TLR monomers in violet and red, MD-2 dimers shown in grey. Lipid A species and side chains are shown in licorice representation (cyan – carbon, red – oxygen, blue – nitrogen, brown – phosphorus). All values in panels (B)-(D) and (F) were averaged over last 200 ns and across both dimers.

Ligand-induced conformational changes that bring the C-terminal regions of TLRs together are important for the activation of downstream signalling. We then hypothesized that the loss of receptor stability in *m/z* 1,388 systems may lead to separation of the membrane-proximal C-terminal regions of TLR4. To test this, we performed principal component analysis (PCA) for the TLR4 Cα atoms, in order to filter the “noise” from the trajectory and identify the dominant collective motions of the receptor chains. In simulations of the active TLR4_2_/MD-2_2_ complex *E. coli* lipid A revealed no significant “separating” motions of TLR4 chains, consistent with stabilization of the complex when bound to agonist (Fig 4I). Similar results were obtained testing lipid A species *m/z* 2,007*, m/z* 1,797 and *m/z* 1,824 (Fig 4I). In contrast, in the case of *m/z* 1,388 we detected vertical sliding motion between the two chains (Fig 4I). This pattern of motion is consistent with a loss of proximity of the membrane-proximal C-terminal regions and inactivation of the receptor complex.

Collectively, the MD simulations demonstrate that the hexa-acylated and hepta-acylated lipid A species found in pathogenic and non-pathogenic *Yersiniae* lead to stabilization of the TLR4_2_/MD-2_2_ receptor complex in a similar way as the benchmark *E. coli* lipid A and, therefore, it is expected that they are agonists of TLR4. On the contrary, the presence of the tetra-acylated lipid A species *m/z* 1,388 found only in pathogenic *Yersiniae* grown at 37°C within the active TLR4_2_/MD-2_2_ receptor complex leads to destabilization of the complex, supporting the notion that this lipid A species is a poor agonist of TLR4.

### TLR4 recognition of the lipid A PAMP

We next sought to provide experimental evidence to support the results obtained by MD simulations suggesting differences in the activation of TLR4 between hexa-acylated and hepta-acylated lipid A species on the one hand, and the tetra-acylated ones on the other hand. Ligand-dependent dimerization of TLR4/MD-2 heterodimers results in endocytosis of the active TLR4_2_/MD-2_2_ complex [36–38]. Therefore, the loss of TLR4 from the plasma membrane represents the most proximal event in the initiation of TLR4 pathway activation. This event can be monitored by flow cytometry and represents a sensitive quantitative assay to assess the interaction of the lipid A PAMP with endogenous PRRs in the natural cellular context (the macrophage surface [36–38]. Live *E. coli* expressing hexa-acylated lipid A interacts efficiently with TLR4 [36–38]; and, therefore, it was used as a benchmark to delineate the engagement of different *Yersinia* strains expressing different lipid A PAMPs with TLR4 as we did for the MD simulations. Pathogenic, YeO8 and YeO3 strains grown at 21°C behaved similarly to *E. coli*, in that they triggered the loss of TLR4 (Fig 5A). Of note, these lipid As present modifications with aminoarabbnose and palmitate (Table 1 and Table 2), suggesting that the decorations of the lipid A PAMP had no effect on the internalization of TLR4. These infections were done with the pYV-cured strains to avoid any effect due to the activation of the YsC T3SS upon contact with macrophages [39–41]. Non-pathogenic strains, *Y. aldovae*, *Y. nurmii* and *Y. enterocolitica* 1A 0902, grown at 21°C also induced the loss of TLR4 from the membrane (Fig 5A). These lipid A s are also decorated with aminoarabinose and palmitate (Table 1 and Table 2), providing further evidence that the decorations do not affect the engagement with TLR4. In sharp contrast, infection with pathogenic strains grown at 37°C did not result in the internalization of the receptor (Fig 5A) whereas non-pathogenic strains engaged with TLR4 (Fig 5A). This is so even in the case of *Y. aldovae* lipid A that it is decorated with palmitate (Table 1 and Table 2). No significant differences were observed on the engagement with TLR4 between non-pathogenic strains grown at 21°C and at 37°C (p > 0.05 for each comparison; Fig 5A). Altogether, this analysis established that the deacylated lipid A PAMP expressed by pathogenic strains at 37°C limits engagement with TLR4. The data also support the notion that the lipid A decorations with aminoarabinose and palmitate do not affect engagement with TLR4.

**Figure 5.**
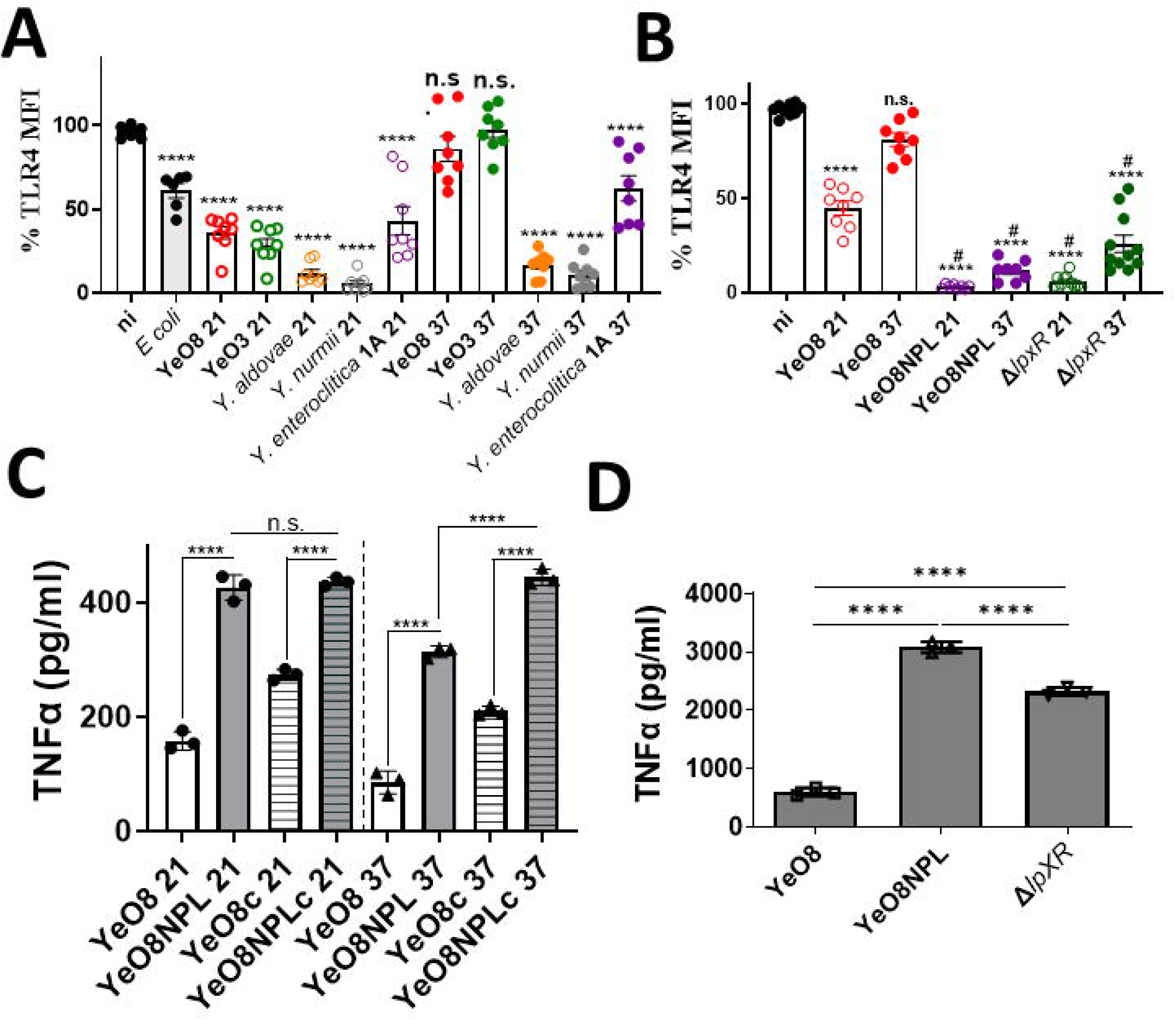
Engagement of Yersinia lipid As with TLR4 and subsequent inflammation. (A) Surface expression of TLR4 as measured by mean fluorescence intensity (MFI) on iBMDMs non-infected (ni) or exposed to live *E. coli*, pathogenic *yersiniae,* YeO8 and YeO3, and non- pathogenic *yersiniae*, 1A strain 0902, *Y. aldovae, Y. nurmii*, grown at 21°C (denoted as 21) and 37°C (denoted as 37). (B)) Surface expression of TLR4 as measured by MFI on iBMDMs non-infected (ni) or exposed to live YeO8, chimeric YeO8 strain expressing a non-pathogenic lipid A, strain YeO8NPL, and YeO8 *lpxR* mutant (Δ*lpxR*, strain YeO8-Δ*lpxR*) grown at 21°C (denoted as 21) and 37°C (denoted as 37). (C) TNFα secretion by infected iBMDMs with YeO8,and YeO8NPL.“c” denotes bacteria without the virulence plasmid. Strains were grown at 21°C (denoted as 21) and 37°C (denoted as 37). (D) TNFα secretion by iBMDMs challenge with 10 ng/ml of repurified LPS from YeO8, YeO8NPL, and YeO8 *lpxR* mutant (Δ*lpxR*, strain YeO8-Δ*lpxR*) grown at 37°C. Data are presented as mean ± SD (n□=□3). In panels A and B, \*\*\*\**P*□≤□0.0001; ns, *P*□>□0.05 for the comparisons against non-infected cells, and # *P*□≤□0.0001 for the comparisons against YeO8 grown at 37°C using One□way ANOVA with Bonferroni contrast for multiple comparisons test. In panels C and D, \*\*\*\**P*□≤□0.0001; ns, *P*□>□0.05 for the indicated comparisons using One□way ANOVA Dunnett’s multiple comparisons test.

We next asked whether changes in the lipid A PAMP expressed by pathogenic *Yersiniae* would be sufficient to affect the interaction with TLR4. To address this question, we constructed a pathogenic chimeric strain expressing the lipid A of non-pathogenic *Yersiniae*. First, we constructed a double *lpxR* and *lpxL* mutant in the YeO8 background, and then we introduced *lpxL* from *Y. aldovae* at the single copy in the chromosome using the Tn7 delivery system to generate strain YeO8NPL. We extracted the lipid A from YeO8NPL and analysed its structure by MALDI-TOF. Figure S2 shows that when YeO8NLP was grown at 21°C (Fig S4A), the lipid A contained species *m/z* 1,768, *m/z* 1,797 and *m/z* 1,824 corresponding to hexa-acylated lipid As, and *m/z* 2,063 corresponding to an hepta-acylated lipid A. Species *m/z* 1768 and *m/z* 1,1824 are modified with aminoarabinose, *m/z* 1,899 and *m/z* 1954. When grown at 37°C, lipid A from YeO8NPL contained species *m/z* 17668 and *m/z* 1,797 (Fig S4B). No tetra-acylated lipid A species were found at any temperature. Altogether, these results demonstrate that YeO8NPL expressed the same lipid A as non-pathogenic *Yersiniae*. Infection with YeO8NPL, grown either at 21°C or at 37°C, induced the loss of TLR4 from the membrane (Fig 5B), and the levels were significantly lower than those found in cells infected with YeO8 (Fig 5B). No differences were observed in TLR4 engagement between YeO8NPL grown either at 21°C or at 37°C (p>0.05; Fig 5B). The fact that the levels of TLR4 in cells infected with the *lpxR* mutant grown either at 21°C or at 37°C were significantly lower than those found in cells infected with YeO8 (Fig 5B) further demonstrates the notion that increased acylation of the lipid A PAMP increases the engagement with TLR4.

We reasoned the differences in engagement with TLR4 between *Yersiniae* expressing different lipid A PAMPs should translate to differences in TLR4-dependent inflammatory responses. When we challenged macrophages with bacteria grown at 21°C, YeO8NPL induced higher levels of TNFα than the wild-type strain (Fig 5C). Similar results were observed when macrophages were challenged with bacteria grown at 37°C (Fig 5C) although the levels were lower than those found in cells infected with bacteria grown at 21°C (Fig 5). This was dependent on the well-known anti-inflammatory action of the pYV-encoded Ysc T3SS [42, 43] because the TNFα levels induced by wild-type bacteria cured of the pYV were higher than those induced by bacteria harbouring the virulence plasmid. This was not the case for YO8NPL grown at 21°C because we did not detect any significant difference in the TNFα levels induced by pYV positive and cured bacteria (Fig 5C).

Nevertheless, at 21 and 37°C YeO8NLP cured of the pYV induced higher TNFα levels than the wild-type strains cured of the plasmid (Fig 5C). We next isolated LPS to evaluate whether purified LPS behaved similarly to whole bacteria. LPS purified from YeO8NPL grown at 37°C induced more TNFα than LPS isolated from YeO8 and the *lpxR* mutant (Fig 5D), the levels induced by the latter were higher than those induced by the wild-type LPS (Fig 5D).

Altogether, these results demonstrate that the lipid A acylation pattern dictates the engagement with TLR4 and the activation of inflammatory responses. The lipid A PAMP from non-pathogenic bacteria engages efficiently with TLR4 and evokes a higher inflammatory response than that of pathogenic bacteria. Switching the lipid A PAMP expressed by pathogenic bacteria to a non-pathogenic PAMP is sufficient to modify the recognition by TLR4 and subsequent activation of inflammation.

### Interaction of the lipid A PAMP with antimicrobial peptides

Likewise PRRs, antimicrobial peptides interact with the lipid A PAMP [44, 45]. The decorations of the lipid A counteract this binding and, therefore, they protect Gram-negative pathogens against antimicrobial peptides [46]. The temperature-dependent variations in the modifications of the lipid A from pathogenic and non-pathogenic *Yersiniae* led us to test the susceptibility to antimicrobial peptides of the *Yersinia* strain. We and others have used polymyxin B and magainin II to probe the contribution of the modification of the lipid A with aminoarabinose and palmitate, respectively, to bacterial susceptibility to antimicrobial peptides [23, 47–49]. When bacteria were challenged with polymyxin B, we observed that bacteria grown at 21°C were significantly more resistant than those grown at 37°C (Fig 6A). These results are consistent with the detection of lipid A modified with aminoarabinose in the lipid As produced by *yersiniae* grown only at 21°C. We and others have previously shown that the modification of the lipid A with aminoarabinose mediates resistance to polymyxin B in *Yersiniae* [23–25]. No significant differences were found between pathogenic and non-pathogenic strains at any growth temperature (Fig 6A). YeO8, YeO3 and YeO9 were also more susceptible to magainin II when grown at 37°C than at 21°C (Fig 6B). This is consistent with the lack of lipid A modification with palmitate at 37°C (Fig 1). Previous work demonstrated that the lipid A modification with palmitate mediates resistance to magainin II [23, 48, 49]. In the case of the non-pathogenic strains, the susceptibility to magainin II differentiated two groups. Those strains more susceptible at 37°C than 21°C like the pathogenic strains, and those showing no significant differences in the susceptibility between temperatures (Fig 6B). The latter group includes *Y. bercovieri, Y. mollaretii, Y. frederiksenii, Y. intermedia, Y. aldovae*, and *Y. kristensenii*; these strains present the modification of the lipid A with palmitate at both growth temperatures (Table S2). Altogether, these results indicate that the resistance to antimicrobial peptides is not a unique trait of pathogenic bacteria and, therefore, the modifications of the lipid A PAMP leading to resistance cannot be strictly considered a result of the host-pathogen co-evolution to evade their antimicrobial action.

**Figure 6.**
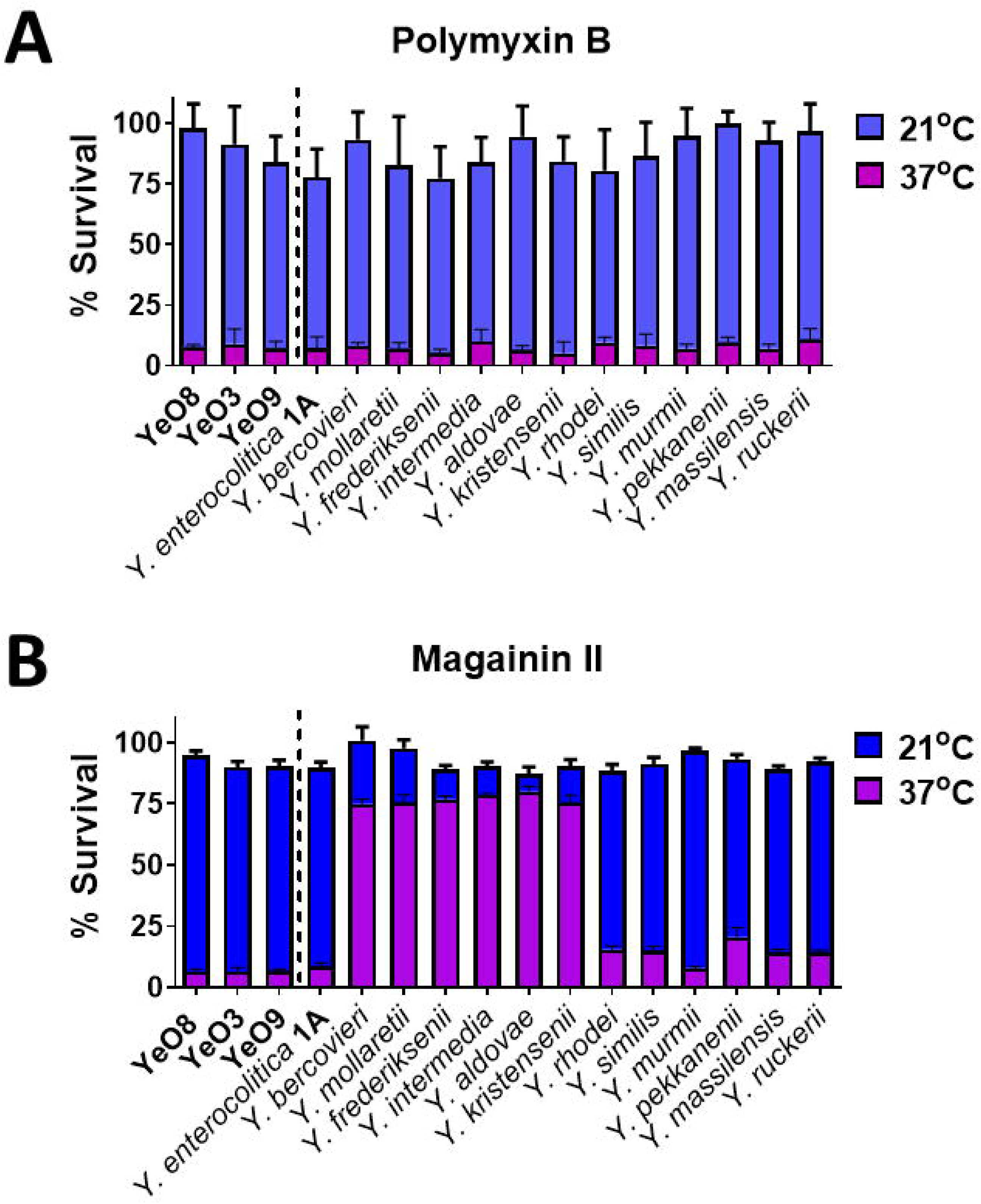
Susceptibility of *Yersiniae* strains to antimicrobial peptides. (A) Susceptibility of pathogenic and non-pathogenic *Yersinia*e strains grown at 21oC or 37oC to polymyxin B (10 µg/ml). (B) Susceptibility of pathogenic and non-pathogenic *Yersiniae* strains grown at 21oC or 37oC to magainin II (10 µg/ml). The data are presented as mean ± SD (n□=□3).

### Effect of the lipid A PAMP on the function of *Yersinia* virulence factors

The ability to switch the lipid A PAMP of pathogenic *Yersinia* to a non-pathogenic lipid A PAMP offered the unique opportunity to ascertain the effect of the lipid A PAMP from non-pathogenic *yersiniae* on the expression and function of *Yersinia* virulence factors. We investigated the flagellar regulon, invasin (Inv), responsible for the for the invasion of the host, and the anti-host Ysc T3SS, encoded in the pYV virulence plasmid.

#### (i) Flagellar regulon and the lipid A PAMP

The flagellar regulon regulates several virulence genes and, therefore, contributes to *Y. enterocolitica* pathogenesis [50]. Moreover, motility is important for the invasion of cells [51]. In vitro, YeO8 is motile when grown at 21°C but not at 37°C [52, 53]. To examine the influence of the non-pathogenic lipid A PAMP on the flagellar regulon, we first quantified the migration of the strains in motility medium (1% tryptone-0.3% agar plates). YeO8NPL was more motile than the wild-type strain (Fig 7A and 7B). In contrast, and corroborating previous results [20], the *lpxR* mutant was less motile than the wild-type (Fig 7B), suggesting that the increased acylation of YeO8NPL is not responsible for the enhanced motility. On the other hand, the expression of *Y. aldovae lpxL* in the background of the YeO8 *lpxL* mutant was sufficient to increase the motility (Fig 7B). YeO8 *lpxL* did not affect the motility of the *lpxL* mutant (Fig 7B). We have previously shown that absence of *lpxL* does not affect the motility of YeO8 [21]. These results indicate that the expression of *Y. aldovae lpxL* increases the motility of pathogenic *yersiniae*.

**Figure 7.**
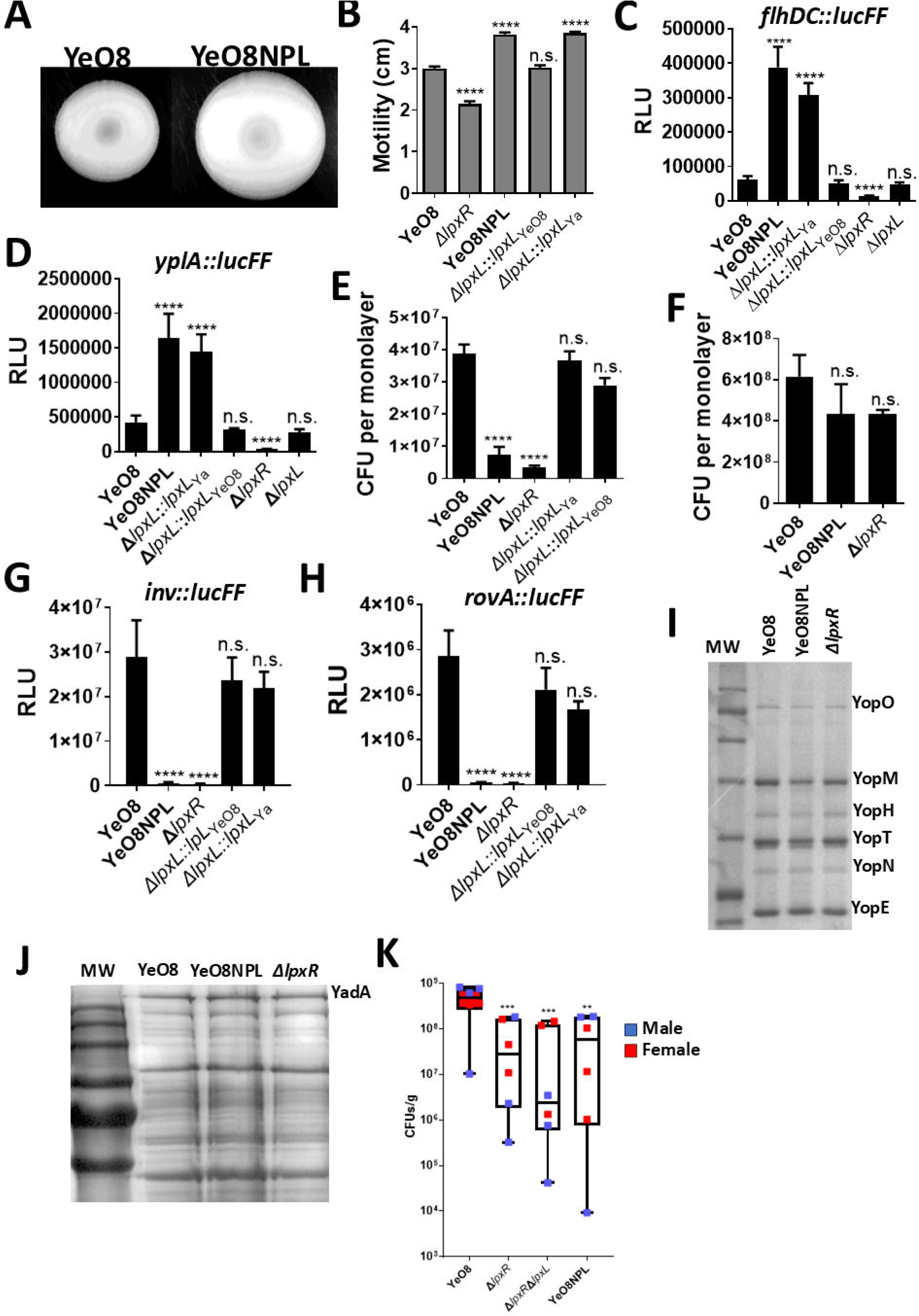
Effect of the lipid A PAMP on *Yersinia* virulence. (A) Motility assay with YeO8 and YeO8NPL in a semisolid agar plate (0.3% agar and 1% tryptone). Plates were incubated at 22°C for 24 h. (B) Quantification of the diffusion diameter of YeO8, and YeO8 *lpxR* mutant (Δ*lpxR*, strain YeO8-Δ*lpxR*), YeO8 *lpxL* mutant complemented with YeO8 *lpxL* (Δ*lpxL::lpxL_YeO8_*) and YeO8 *lpxL* mutant complemented with *Y.aldovae lpxL* (Δ*lpxL::lpxL_Ya_*,). (C) Analysis of *flhDC* expression by YeO8, YeO8NPL, YeO8 *lpxL* mutant complemented with YeO8 *lpxL* (Δ*lpxL::lpxL_YeO8_*), YeO8 *lpxL* mutant complemented with *Y.aldovae lpxL* (Δ*lpxL::lpxL_Ya_*,)YeO8 *lpxR* mutant (Δ*lpxR*, strain YeO8-Δ*lpxR*), and YeO8 *lpxL* mutant (Δ*lpxL*, strain YeO8-Δ*lpxL)* carrying the transcriptional fusion *flhDC::lucFF* grown at 21°C. (D) Analysis of *yplA* expression by YeO8, YeO8NPL, YeO8 *lpxL* mutant complemented with YeO8 *lpxL* (Δ*lpxL::lpxL_YeO8_*), YeO8 *lpxL* mutant complemented with *Y.aldovae lpxL* (Δ*lpxL::lpxL_Ya_*,), YeO8 *lpxR* mutant (Δ*lpxR*, strain YeO8-Δ*lpxR*), and YeO8 *lpxL* mutant (Δ*lpxL*, strain YeO8-Δ*lpxL)* carrying the transcriptional fusion *yplA::lucFF* grown at 21°C.(E) Invasion of HeLa cells by YeO8, YeO8NPL, YeO8 *lpxL* mutant complemented with YeO8 *lpxL* (Δ*lpxL::lpxL_YeO8_*), YeO8 *lpxL* mutant complemented with *Y.aldovae lpxL* (Δ*lpxL::lpxL_Ya_*,), YeO8 *lpxR* mutant (Δ*lpxR*, strain YeO8-Δ*lpxR*). Invasion assays were done in triplicate without centrifugation (n□=□3).(F) Adhesion to HeLa cells by YeO8, YeO8NPL, and YeO8 *lpxR* mutant (Δ*lpxR*, strain YeO8-Δ*lpxR*). Adhesion assays were done in triplicate without centrifugation (n□=□3) (G) Analysis of *inv* expression by YeO8, YeO8NPL, YeO8 *lpxL* mutant complemented with YeO8 *lpxL* (Δ*lpxL::lpxL_YeO8_*), YeO8 *lpxL* mutant complemented with *Y.aldovae lpxL* (Δ*lpxL::lpxL_Ya_*,), and YeO8 *lpxR* mutant (Δ*lpxR*, strain YeO8-Δ*lpxR*) carrying the transcriptional fusion *inv::lucFF* grown at 21°C. (H) Analysis of *rovA* expression by YeO8, YeO8NPL, YeO8 *lpxL* mutant complemented with YeO8 *lpxL* (Δ*lpxL::lpxL_YeO8_*), YeO8 *lpxL* mutant complemented with *Y.aldovae lpxL* (Δ*lpxL::lpxL_Ya_*,), and YeO8 *lpxR* mutant (Δ*lpxR*, strain YeO8-Δ*lpxR*) carrying the transcriptional fusion *rovA::lucFF* grown at 21°C. (I) SDS-PAGE (the acrylamide concentration was 4% in the stacking gel and 12% in the separation one) and Coomasie brilliant blue staining of proteins from the supernatants of Ca^2+^- deprived cultures from YeO8, YO8NPL and YeO8 *lpxR* mutant (Δ*lpxR*, strain YeO8-Δ*lpxR*). Result is representative of three independent experiments. (J) SDS-PAGE (the acrylamide concentration was 4% in the stacking gel and 10% in the separation one) followed by Coomasie brilliant blue staining of cell extracts from YeO8, YO8NPL and YeO8 *lpxR* mutant (Δ*lpxR*, strain YeO8-Δ*lpxR*) grown in RPMI 1640 at 37°C. YadA protein is marked. Result is representative of three independent experiments. (K) CFUs per gram of Peyer’s patches from male and female mice infected with YeO8, YeO8NPL, YeO8 *lpxR* mutant (Δ*lpxR*, strain YeO8-Δ*lpxR*) and YeO8 *lpxR-lpxL* double mutant (Δ*lpxR* Δ*lpxL*, strain YeO8-Δ*lpxR-* Δ*lpxL*). Data are presented as mean ± SD (n□=□3). In all panels, \*\*\*\**P*□≤□0.0001; \*\*\**P*□≤□0.001; \*\**P*□≤□0.01; ns, *P*□>□0.05 for the comparisons against non-infected cells, and # *P*□≤□0.0001 for the comparisons against YeO8 using One□way ANOVA with Dunnett’s multiple comparisons test.

*Yersinia* motility is related to the levels of flagellins which, in turn, are regulated by the expression of *flhDC*, the flagellum master regulatory operon [52]. We then hypothesized that the expression of *flhDC* could be higher in YeO8NPL due to the expression of *Y. aldovae* LpxL. To test this hypothesis, the *flhDC::lucFF* transcriptional fusion [53] was introduced into the chromosome of the strains and the luciferase activity was determined. Validating our hypothesis, luminescence was higher in the YeO8NPL and YeO8 *lpxL* mutant complemented with *Y. aldovae lpxL* than in the wild type strain (Fig 7C). No differences were observed in the luciferase levels between the wild-type strain and the *lpxL* mutant (Fig 7C). Further supporting previous results [20], luminiscence was lower in the *lpxR* mutant than in the wild type, and there were no differences in luminescence between the YeO8, the *lpxL* mutant, and the complemented strain expression YeO8 *lpxL* (Fig 7C). To corroborate further that the flagellar regulon is upregulated in YeO8NPL, we next assessed the expression of the phospholipase *yplA* whose expression is regulated by *flhDC* [50, 54]. The luciferase activity of the transcriptional fusion *yplA::lucFF* was higher in the YeO8NPL and YeO8 *lpxL* mutant complemented with *Y. aldovae lpxL* backgrounds than in the wild type, the *lpxL* mutant and the mutant complemented with YeO8 *lpxL* (Fig 7D). No significant differences were observed between the later strains (Fig 7D). In contrast, the luciferase levels were lower in the *lpxR* mutant than in the wild-type strain (Fig 7D). Altogether, these results demonstrate that the flagellar regulon is upregulated in pathogenic *yersiniae* expressing the non-pathogenic *yersiniae* lipid A PAMP triggered by the expression of *Y. aldovae lpxL*.

#### (ii) Invasin and the lipid A PAMP

Inv is an outer membrane protein responsible for the invasion of the host [55, 56]. We then asked whether the invasion of cells is affected by the lipid A PAMP using a gentamicin protection assay. The number of intracellular bacteria in HeLa cells was 80% lower in cells infected with YeO8NPL and the *lpxR* mutant (Fig 7E) whereas no differences were found between cells infected with the wild-type strain and the *lpxL* mutant complemented with *Y. aldovae lpxL* or YeO8 *lpxL* (Fig 7E). The differences in invasion are not due to differences in attachment to the cells because the number of bacteria attached to HeLa cells was not significantly different between cells infected with the wild type, YeO8NPL or the *lpxR* mutant (Fig 7F). These differences in the invasion of cells led us to assess the expression of *inv*. We constructed an *inv::lucFF* transcriptional fusion which was introduced into the different strains, and *inv* expression was monitored as luciferase activity. Luminescence was lower in the YeO8NPL and the *lpxR* mutant backgrounds than in the wild-type strains and the *lpxL* mutant complemented with YeO8 *lpxL* or *Y. aldovae lpxL* (Fig 7G), suggesting that the reduced expression of *inv* mediates the differences in cell invasion between strains.

RovA regulates the expression of *inv* [57, 58]. Therefore, we sought to determine whether the low *inv* expression correlates with the expression of *rovA*. A *rovA::lucFF* transcriptional fusion [21] was introduced in to the strains and the luciferase activity determined. The expression of *rovA* was also downregulated in YeO8NPL and the *lpxR* mutant backgrounds (Fig 7H). No differences were found between the wild type and the *lpxL* mutant complemented with YeO8 *lpxL* or *Y. aldovae lpxL* (Fig 7H). Collectively, this evidence establishes that the increase acylation of the lipid A in YeO8 expressing the non-pathogenic lipid A PAMP results in a decrease expression of *inv* and its positive regulator *rovA* with a concomitant decrease in the invasion of cells.

#### (iii) pYV-encoded virulence factors and the lipid A PAMP

The Ysc T3SS, encoded in the pYV virulence plasmids, is required for *Yersinia* virulence and it secretes a set of protein effectors called Yops that enable *Y. enterocolitica* to multiply extracellularly in lymphoid tissues [15]. Analysis of Yop secretion showed no differences between the wild-type, the *lpxR* mutant, and YeO8NPL (Fig 7I). The disruption of the cytoskeleton upon injection of YopE to cells is one of the most sensitive read-outs to assess the activity of the Ysc T3SS [59]. YeO8, YeO8NPL and the *lpxR* mutant induced similar disruption and condensation of the actin microfilament structure whereas this was not the case in cells infected with the *yopE* mutant (Fig S5A and Fig S5B).

YadA is another pYV-encoded factor mediating bacterial adhesion, bacterial binding to proteins of the extracellular matrix and complement resistance [60, 61]. We analysed the expression of the YadA trimeric form by SDS-PAGE followed by Coomasie blue staining. Figure 7J shows that YeO8, YeO8NPL and the *lpxR* mutant produced the same amount of YadA. To investigate YadA functionally, we assessed the binding of YadA expressing bacteria to collagen using as negative control YeO8 cured of the pYV. We observed no differences in collagen binding between YeO8 and YeO8NPL (Fig S5C and Fig S5D) whereas, as expected, the strain cured of the pYV did not bind to collagen (Fig S5C and Fig S5D). We have demonstrated that the *lpxR* mutant binds to collagen as well as YeO8 [20].

Taken together, these results suggest that expression of the non-pathogenic lipid A PAMP does not affect the production and function of pYV-encoded virulence factors.

### Effect of the lipid A PAMP on *Yersinia* virulence

To establish the effect on virulence of switching the lipid A PAMP, BALB/c mice were infected orogastrically, and 2 days postinfection, mice were euthanized, and the. numbers of bacteria present in the Peyer’s patches were determined by plating. All strains colonized the Peyer’s patches (Fig 7K). However, the loads of the *lpxR* mutant were seven times lower than those of the wild-type strain (Fig 7K). The double mutant lacking *lpxR* and *lpxL* was even more impaired than the *lpxR* mutant to colonize the Peyer’s patches; the bacterial loads were eleven and two times lower than those of the wild type and *lpxR* mutant, respectively (Fig 7K). The bacterial loads of YeO8NPL were not significantly different than those of the double mutant (p> 0.05) and six times lower than that of the wild-type strain, demonstrating that a switch from a pathogenic to a non-pathogenic lipid A PAMP is enough to attenuate virulence.

## DISCUSSION

This study was designed to address one of the fundamental questions of the pattern recognition concept: is there a PAMP associated with pathogenicity shaped by the challenge imposed by the innate immune system? By probing the lipid A PAMP, a key molecular pattern expressed by Gram-negative bacteria, we drew three conclusions. First, we demonstrated that the reduced acylation of the lipid A PAMP is an evolved adaptation to evade recognition by mammalian LPS receptors, limiting the activation of inflammatory protective responses and, therefore, contributing to virulence. Second, the decorations of the lipid A PAMP do not represent a unique adaptation to a mammalian ecosystem and are shared by pathogenic and non-pathogenic bacteria. And third, virulence factors cannot overcome a switch in the lipid A PAMP associated with pathogenicity, resulting in decrease virulence.

By using MD simulations and a highly sensitive flow cytometry assay to detect endogenous TLR4, we have demonstrated that the reduced acylation of the lipid A PAMP results in reduced interaction with TLR4 with a concomitant decrease in inflammation. Our work provides a conceptual framework to explain why so many pathogens including *Pseudomonas aeruginosa*, *Helicobacter pylori*, *Francisella turalensis*, *Porphyromonas gingivalis*, as well as members of the microbiome such as *Prevotella intermedia* and *Bacteroides spp*, express lipid As with reduced acylation [62–75]. This is also in agreement with evidence demonstrating that the mammalian host environment induces a reduction in the lipid A acylation state not observed in bacteria culture in vitro as it has been demonstrated for *Klebsiella pneumoniae* [76]. Noteworthy, a common feature shared by all these different lipid As is the comparative lower activation of the innate immune system compare to lipid As similar to the so-called canonical “hexa-acyl lipid A”. This correlates with a body of work probing diverse lipid As generated by bacterial glycoengineering in which those lipid As with reduced acylation induce less inflammation [77, 78].

It is interesting to note that the hepta-acylation of the lipid A found in non-pathogenic strains when grown at 37°C, for example *Y. aldovae*, is not associated with reduced engagement with TLR4 and a decrease inflammation as our MD simulations and experimental evidence establishes. This is in sharp contrast to previous reports suggesting that PagP-controlled hepta-acylation results in limited TLR4 activation [79–82]. However, our results are consistent with observations made probing *Acinetobacter baumannii*, *Pseudomonas aeruginosa*, and *E. coli* expressing hepta-acylated lipid A [78, 83, 84]. Likewise our work, these studies assessed LPS-mediated inflammatory responses interrogating live bacteria instead of testing purified LPS, highlighting the importance of interrogating the lipid A PAMP in the physiological context of the outer membrane.

Our results emphasize the role of the Ysc T3SS to overcome inflammation induced by the lipid A PAMP. However, and despite its efficiency, the pYV-encoded T3SS cannot control the inflammation induced by the non-pathogenic lipid PAMP. Overall, these findings support the notion that the low inflammatory response associated to infections by pathogenic *Yersiniae* is dependent on the reduced activation of the LPS receptor complex induced by the pathogenic lipid PAMP couple to the anti-inflammatory action of the Ysc T3SS.

There is a wealth of evidence demonstrating that the decorations of the lipid A PAMP contribute to the virulence of Gram-negative pathogens [46]. Our results do not challenge these findings; however, they demonstrate that this is not a unique trait of pathogenic bacteria. The decorations of the lipid A fortify the outer membrane, acting as a defence mechanism against antimicrobial peptides and other toxins [46]. Undoubtedly, these lipid A structural changes are also useful for bacteria living in the environment where they could be also exposed to these agents produced by other microbes such as *Streptomycetes*, plants, and aquatic and marine organisms, explaining why the lipid A decorations are not unique to bacteria colonizing or infecting mammalian hosts. In the *Yersinia* context, it was previously believed that the temperature-dependent expression of these lipid A decorations represented an adaptation to the environment found in the mammalian gut and lung mucosae [23, 24]. Here, we have shown that this is not the case because non-pathogenic species also display this trait. In pathogenic strains, the temperature-dependent regulation of the loci controlling lipid A modifications is explained by H-NS-dependent negative regulation alleviated by RovA [23]. Interestingly, non-pathogenic *Yersiniae* do not encode *rovA*; however, they do encode *slyA* which has been shown to alleviate the negative regulation exerted by H-NS in other Gram- negative pathogens [85–87]. We posit the SlyA-H-NS circuit controls the expression of the loci controlling the lipid A decorations in non-pathogenic *Yersiniae*. Future studies are warranted to validate this notion.

One striking finding of our study is the dramatic effect on virulence of switching the lipid A PAMP. Increasing the acylation pattern, as occurs in the *lpxR* mutant, already resulted in an attenuation of virulence. Remarkably, expressing the non-pathogenic lipid A PAMP triggered an even further attenuation. This finding cannot be explained by lack of function of the Ysc T3SS because we did not observe any differences in the production and function of the pYV-encoded Yops and YadA between the wild-type strain, YeO8NPL and the *lpxR* mutant. At least two non-exclusive explanations may account for the low-level colonization of Peyer’s patches: (i) switching the lipid A PAMP results in a decrease penetration of the intestinal epithelium, and (ii) YeO8NPL is cleared by a host defence mechanism present in the tissue. In support of the former, *inv* expression was lower in YeO8NPL than in the wild type and the mutant was impaired in its ability to invade epithelial cells. However, the reduced *inv* expression cannot explain fully the low bacterial loads in the Peyer’s patches of mice infected with YeO8NPL because no differences in *inv* expression were found between the *lpxR* mutant and YeO8NPL. Supporting the latter, the heightened inflammation triggered by YeO8NPL upon TLR4 recognition of the lipid A PAMP will result undoubtedly in vivo in the activation of host protective proinflammatory responses. This agrees with previous studies illustrating the crucial role of TLR4 signalling to control *Yersinia* infections [88–90].

## MATERIALS AND METHODS

### Ethics

Balb/c mice were purchased from Charles River. Mice were age and sex-matched and used between 8-12 weeks of age. The experiments involving mice were approved by the Queen’s University Belfast’s Ethics Committee and conducted in accordance with the UK Home Office regulations (project licence PPL2910) issued by the UK Home Office. Animals were randomized for interventions but researches processing the samples and analysing the data were aware which intervention group corresponded to which cohort of animals.

### Bacteria strains and growth conditions

Bacterial strains are described in Table S2. Bacteria were grown overnight in 5 mL of Miller’s Luria Broth (LB) medium (Melford) at 37°C or 21°C on an orbital shaker (180 rpm). Overnight bacterial cultures were refreshed 1/10 into a new tube containing 4.5 mL of fresh LB. After 3 h at 37°C, bacteria were pelleted (2,500x g, 20 min, 22°C), and resuspended in PBS to an OD_540_ of 0.8 (corresponding to 4×10^8^ CFUs/ml). Where appropriate, antibiotics were added to the growth medium at the following concentration: ampicillin (Amp), 50 μg/ml for *E. coli* and *Yersinia spp*, and 100 μg/ml for *Yersinia spp* in agar plates; kanamycin (Km), 50 μg/ml for *E. coli* and *Yersinia spp*, and 100 μg/ml for *Yersinia spp* in agar plates; gentamycin (Gm), 100 μg/ml; chloramphenicol (Cm), 25 μg/ml; tetracycline (Tet), 12.5 μg/ml; streptomycin (Str), 100 μg/ml; trimethroprim (Tmp), 100 μg/ml.

To cure the pYV plasmid from pathogenic *Yersinia*, bacteria were grown at 37°C in Congo Red Magnesium oxalate agar plates [91]. Colony size and lack of uptake of Congo Red were used to detect loss of the virulence plasmid. This was further confirmed by testing the YadA-dependent autoagglutination ability [92].

### Isolation of lipid A and MALDI-TOF analysis

Bacterial cultures were grown overnight at 21 °C or 37 °C in a 5 ml of LB. On the next day, the cultures were used to inoculate 1 litre Erlenmeyer containing 250 ml of LB broth (cultured at the same temperature and agitation). After 24 hours, the bacterial cells were harvested by centrifugation (2,000 x g, 15 minutes, room temperature), washed twice in phosphate buffer (PB, 1.5 g of Na_2_HPO_4_, 0.2 g of KH_2_PO_4_). The final pellet was resuspended in 10 ml of PB and placed at -80°C. The cell pellet was lyophilized.

The lipid A extraction was carried out by an ammonium hydroxide / isobutyric acid method [21, 93]. Briefly, 10 mg of the lyophilized bacterial was resuspended in 400 μl of a solution of butiric acid (98% pure Sigma-Aldrich) / 1M ammonium hydroxide (5:3, v/v), and incubated during 2 hours at 100°C with vortexing every 15 minutes. Then, the hydrolysed product was cooled down (5 minutes incubation in ice), and centrifuged (2,000 x g, 15 minutes, room temperature). The supernatant was then transferred to a new microcentrifuge tube and the same volume of water was added (1:1, v/v). Next, the samples were frozen at -80°C and lyophilized. After, the samples were washed in 400 μl of methanol twice and centrifuged (2,000 x g, 15 minutes, room temperature). The final pellet was solubilized in a mixture of chlorophorm/methanol/water (3:1.5:0.25, v:v:v), as a final concentration 1mg/ml. Each sample was treated with traces of the ion-exchange resin (Dowex 50W-X8; H+). 1 μl of the lipid A suspension were spotted on a MALDI-TOF target, dried (5 min at room temp), and 1 μl of matrix was added on top. The 2,5-Dihydroxybenzoic acid (Bruker) was used as a matrix dissolved in (1:2) acetonitrile-0.1% trifluoroacetic acid (Following manufactureŕs instructions). The lipid A analysis was performed by mass spectrometry using a Bruker autoflex speed TOF/TOF mass spectrometer (Bruker Daltonics Inc.) in negative reflective mode. Each spectrum was obtained as an average of at least 1000 shots (ion-accelerating voltage set at 40kV). Two controls were used for calibration: Peptide Calibration Standard II (Bruker), a mixture of known peptide sizes, and lipid A extracted from the *E.coli* strain MG1655 grown in LB at 37 °C. The resulting spectra interpretation was based on previous studies showing that a mass/charge (m/z) above 1000 is proportional to the corresponding lipid A specie.

Construction *Y. aldovae* strain encoding *lpxR*.

We used the Tn7 single copy-based system [28]. The Tn*7* transposon integrates at the site-specific *att*Tn*7*, located downstream of the conserved *glmS* gene, which encodes an essential glucosamine-fructose-6-phosphate aminotransferase [28]*. lpxR* from YeO8 was amplified by PCR using the primers listed in Table S3 and the following cycling conditions: 30 x cycles of: 95°C for 45 sec, 56°C for 45 sec, 72°C for 1.30 min, using 50 ng of genomic DNA, 0.2 mM dNTPs, 10 μM of primers, and 0.2 μl of Pfu DNA polymerase in a final volume of 50 μl). The PCR product was gel-purified (Qiagen Gel extraction kit), phosphorylated, and cloned into PvuII-digested, Antarctic phosphatase treated, and gel-purified, pUC18R6KTmini-Tn7TKm to give pUC18R6KTmini-Tn7TKmLpxR. This vector was mobilized into *Y. aldovae* by triparentral conjugation with the help of a conjugative *E. coli* β2613 carrying the plasmid pTNS2, encoding the Tn7 transposase. The colonies grown were thereafter screened for resistance to kanamycin and susceptibility to ampicillin. Correct integration of the Tn7 transposon was confirmed by PCR using the primers YeO8_GlmS_Tn7comF and Tn7com_R, and Tn7com_F, YeO8_PstS_Tn7comR (Table S3). Additionally, the presence of the lpxR gene was PCR-confirmed using YeO8_lpxR_F, YeO8_lpxR_R primers.

### Complementation of *E. coli* strains

*E. coli* BN1 is an *lpxT, eptA* and *pagP* mutant, producing hexa-acylated lipid A [31]. We previously constructed a *lpxL* mutant of BN1 [30]. This strain was complemented with *lpxL* from YeO8, *Y. aldovae, Y. mollaretti*, and *Y. bercovieri*. The genes were amplified by PCR and cloned into pGEMT-Easy to give pGEMT*lpxL*_YeO8_, pGEMT*lpxL_Y. aldovae_*, pGEMT*lpxL_Y. mollaretti_*, and pGEMT*lpxL_Y. bercovieri_*. The plasmids were introduced into *E. coli* BN1Δ*lpxL* by electroporation. *E. coli* cells were made competent utilizing a microcentrifuge-based procedure (**Choi 2006).

E. coli BN2 is an *lpxT, eptA*, *pagP*, *lpxM* mutant producing penta-acylated lipid A [31]. This strain was complemented with *lpxM* from *Y. aldovae, Y. intermedia*, and *Y. bercovieri*. The genes were amplified by PCR and cloned into pGEMT-Easy to give pGEMT*lpxM_Y. aldovae_*, pGEMT*lpxM_Y. intermedia_*, and pGEMT*lpxM_Y. bercovieri_*. The plasmids were introduced into *E. coli* BN2 by electroporation.

### Complemention of YeO8 *lpxP* mutant

YeO8-Δ*lpxP*GB mutant [21] was complemented using a Tn7-based system. *lpxP* genes from *Y. aldovae, Y. bercovieri* and *Y. intermedia* were amplified by PCR, gel-purified (Qiagen Gel extraction kit), phosphorylated, and cloned into PvuII-digested, Antarctic phosphatase-treated, and gel-purified, pUC18R6KTmini-Tn7TCm [28] to give pUC18R6KTmini-Tn7TCm*lpxP_Y. aldovae_*, pUC18R6KTmini-Tn7TCm*lpxP_Y. bercovieri_*, and pUC18R6KTmini-Tn7TCm*lpxP_Y. intermedia_*. These vectors were mobilized into the YeO8 *lpxP* mutant by triparental conjugation, and the correct integration of the Tn7 transposon at the *att*Tn*7* site confirmed by PCR. The presence of the complementing genes was PCR-confirmed.

### Site-directed mutagenesis of YeO8 *lpxL*

The site-directed mutagenesis of the *lpxL* gene from YeO8 was performed by PCR, as previously described [94] using as template plasmid pGEMT*lpxL*_YeO8_. For that, the primers pairs (Table S3) were designed with a mismatch in the amino acid leucine, position L146. The obtained PCR products were gel purified, phosphorylated with T4 polynucleotide kinase, ligated, and digested with DpnI to break down any remaining template plasmid. The ligated PCR-product was transformed into chemically competent *E. coli* C600 to obtain pGEMT*lpxL*_YeO8Leu146_. Plasmid DNA was isolated from transformants and the *lpxL* gene was sequenced to confirm the generated mutations and to ensure that no other changes were introduced. pGEMT*lpxL*_YeO8Leu146_ was introduced into *E. coli* BN1-Δ*lpxL* by electroporation.

### Construction YeO8NPL, YeO8 strain producing a non-pathogenic lipid A

To construct a YeO8 strain producing the lipid A of non-pathogenic *yersiniae*, we first constructed a double *lpxR-lpxL* mutant which was then complemented with *Y. aldovae lpxL*.

To construct a *lpxL* mutant, two sets of primers were used to amplify 800-900 bp regions flanking the *lpxL* gene. Gel-purified LpxLUp and LpxLDOWN fragments were annealed at their overlapping BamHI sequence in a PCR machine (8 cycles of 95°C for 45 sec, 55°C for 90 sec, 72°C for 90 sec). The reaction included Takara Ex Taq DNA Polymerase, and 0.2 mM dNTPs. The annealed fragment was amplified by PCR using primers YeO8LpxLF1 and YeO8DOWNR1, gel-purified and cloned into pGEMT-Easy to obtain pGEMTΔ*lpxL*. A kanamycin resistance cassette flanked by FRT recombination sites was obtained as a BamHI fragment from pGEMTFRTKm [49] and it was cloned into BamHI-digested pGEMTΔ*lpxL* to generate pGEMTΔ*lpxL*Km. The 3.5 kb *lpxL::Km* allele was obtained by NotI restriction digestion of pGEMTΔ*lpxL*Km, gel-purified and cloned into NotI-digested, Antarctic phosphatase-treated pJTOOL1 [95] to obtain pJTOOLΔ*lpxL*Km. pJTOOL1 is a suicide vector that carries the defective *pir-*negative origin of replication of R6K, the RK2 origin of transfer, and an Cm resistance marker [95]. It also carries the *sacBR* genes that mediate sucrose sensitivity as a positive selection marker for the excision of the vector after double crossover [95]. pJTOOLΔ*lpxL*Km was introduced into the diaminopimelate (DAP) auxotrophic *E. coli* donor strain β2163 [96] and mobilised into the *lpxR* mutant of YeO8 [20] via conjugation.

Bacteria were diluted and aliquots spread on *Yersinia* selective agar medium plates (Oxoid) supplemented with Cm. Bacteria from 5 individual colonies were pooled and allowed to grow in LB without any antibiotic overnight at RT. Bacterial cultures were serially diluted and aliquots spread in LB without NaCl containing 10% sucrose and plates were incubated at room temperature. The recombinants that survived 10% sucrose were checked for their antibiotic resistance. The appropriate replacement of the wild-type alleles by the mutant ones was confirmed by PCR. The kanamycin cassette was excised by Flp-mediated recombination using plasmid pFLP2Tp [20] [97]. The generated mutant was named YeO8-Δ*lpxR*Δ*lpxL*.

*Y. aldovae lpxL* was amplified by PCR, gel-purified, phosphorylated and cloned into PvuII-digested, Antarctic phosphatase treated, and gel-purified, pUC18R6KTmini-Tn7TKm to give pUC18R6KTmini-Tn7TKm*lpxL*_Y. aldovae_. The plasmid was mobilized into YeO8-Δ*lpxR*Δ*lpxL* by triparental mating to obtain YeO8NPL. The correct integration of the Tn7 transposon was assessed by PCR.

### Autoagglutination analysis

YadA is a pYV-encoded outer membrane protein mediating agglutination when bacteria are grown in minimal medium [92]. Therefore, the presence or absence of pYV in pathogenic *Yersinia* can be determined using an autoagglutination test. 1 ml of RPMI1640 media without pH indicator (Gibco) was inoculated with half colony taken from a fresh LB agar plate grown at room temperature for 48 h. The culture was grown at 37 °C, without agitation for 20 h, and the autoagglutination was visually determined.

### Molecular dynamics simulations

All-atom simulations were performed using the GROMACS 2018.3 simulation package [98] utilizing the CHARMM36m force field with the TIP3P water model [99]. The experimental structure (PDB: 3FXI4) of the (TLR4:MD-2)2 heterotetramer bound to *E. coli* ReLPS was used. Protein charges were assigned according to neutral pH with charged termini. Lipid A structures and topologies: i) m/z 1768, ii) m/z 2007, iii) m/z 1797, iv) m/z 1388 and v) m/z 1824 were built using CHARMM-GUI ligand builder & modeler [100], while vi) *E. coli* lipid A was built using CHARMM-GUI membrane builder. In total, six simulation systems were setup of (TLR4:MD-2) complexes bound to a given lipid A type. Each lipid molecule headgroup was aligned to the experimentally solved ReLPS lipid A headgroup. Each dimeric construct was placed in a dodecahedron box with ∼16 nm box edge. Approximately 90,000 TIP3P water molecules were added to the box and 150 mM NaCl salt whilst neutralizing the overall system charge. Energy minimization was performed using the steepest descent minimization algorithm with a 0.01 nm energy step size. The system was equilibrated in a step wise fashion: i) 1 ns with position restraints on protein alpha carbons and lipid headgroups in the NVT ensemble; ii) 20 ns with position restraints on protein alpha carbons and lipid headgroups in the NPT ensemble; iii) 50 ns with position restraints on protein alpha carbons in the NPT ensemble. All position-restrained simulations were run with a force constant of 1,000 kJ mol-1 nm-2. Unrestrained production runs were set to 1000 ns in the NPT ensemble. A temperature of 310 K was maintained using the velocity rescaling thermostat with an additional stochastic term using a time constant of 1 ps. Pressure was maintained semi-isotropically at 1 atm using the Parrinello-Rahman barostat [101] and a time constant of 5 ps. All bonds which involved hydrogens were constrained using the LINCS algorithm [102]. Equations of motion were integrated using the leap-frog algorithm with a time step of 2 fs. Long-range electrostatic interactions were described using the particle mesh Ewald method (PME). The short-range electrostatics reals pace cut-off was 1.2 nm and the short-range van der Waals cut-off was also 1.2 nm. Periodic boundaries conditions were applied in all directions. Simulations were performed on: i) an in-house Linux cluster composed of 8 nodes containing 2 GPUs (Nvidia GeForce RTX 2080 Ti) and 24 CPUs (Intel® Xeon® Gold 5118 CPU @ 2.3 GHz) each as well as on ii) the National Supercomputing Center (https://www.nscc.sg) using 4 nodes containing 128 cores and 256 logical cores (AMD EPYC 7713 @ 2.0 GHz) each.

All simulations snapshots were generated using VMD [103]. The root mean square deviation (RMSD) of the TLR4 dimer backbone was calculated with respect to the experimental structure after alignment onto the same set of atoms. The RMSD of the loop encompassing backbone atoms of residues 120-129 was calculated with respect to the experimental structure after alignment onto each MD-2 monomer backbone. The (TLR4:MD-2)_2_-lipid A contacts were calculated based on a 0.4 nm cutoff distance. The buried solvent accessible surface area (SASA) between two MD-2-lipid and TLR4 dimer was calculated as a sum of MD2-lipid and TLR4 SASA before subtracting (TLR4:MD-2)2–lipid SASA and dividing by two. The RMSDs, protein-lipid hydrogen bonds, protein-lipid contacts and SASA values were calculated as a mean over two monomers and averaged over last 200 ns of trajectory. Principal component analysis was performed on TLR4 dimer alpha carbons. Porcupine plots were generated using extreme conformation of the most dominant principal component (PC1) which corresponded to ∼30% of the total variance captured.

### Modelling LpxL-lipid interactions

YeO8 LpxL sequence was used as bait with the Basic Local Alignment Search Tool (BLAST) at NCBI (http://blast.ncbi.nlm.nih.gov/) to search the non-redundant protein sequence database for similar sequences in the *Yersinia* genus, as well as in *E. coli* (YeO8: A1JN09; *Y. enterocolitica* O3: gi|491299878; *Y. enterocolitica* O9: gi|325666359; *Y. enterocolitica* 1A: gi|571261928; *Y. bercovieri*: gi|238715289; *Y. mollaretii*: gi|238720151; *Y. frederiksenii*: gi|238722579; *Y. intermedia*: gi|238728988; *Y. aldovae*: gi|238704192; *Y. kristensenii*: gi|238699514; *Y. ruckerii*: gi|238707399; *Y. rohdei*: gi|238712349; *Y. similis*: gi|588288876; *Y. nurmii*: gi|902505539; *Y. pekkanenii*: gi|902541928; *Y. massiliensis*: gi|518039378; *E. coli*: P0ACV0). Protein Data Bank (PBD) was searched for crystal structures with bound ligands, which could serve as template for modeling. The only significant hit was LpxM from *A. baumannii* in complex with n-dodecyl-β-D-maltoside and glycerol at 1.99 Å resolution and an E-value of 3e-12 (PDB ID 5KN7) [104], which belongs to the same LPLATs superfamily as LpxL. Since we focused on the conserved hydrophobic lipid binding site of LpxL and the crystal structures deposited in the PDB in general agree much more closely with experimental data where the predicted AlphaFold models and deposited structures differ [105], we did not use ligand free AlphaFold models in this study aiming to predict C12 and C14 binding to LpxL. MALIGN [106] in the BODIL modeling environment [107] was used to align the LpxL sequences from the *Yersinia* genus and *E.coli*, after which the LpxM sequence was added to the alignment using prealigned sequences. For modeling, all sequences except YeO8 LpxL, *Y. aldovae* LpxL or *E. coli* LpxL and the *A. baumannii* LpxM template were deleted from the alignment. A set of 10 models was created with MODELLER [108] and the model with the lowest value of the MODELLER objective function was analyzed and compared with the crystal structure of LpxM. The rotamers of the hydrophobic tunnel were chosen as close to the rotamers in LpxM as possible using Maestro. C12 and C14 were sketched in Maestro and minimized and prepared for docking with LigPrep and force field OPLS4. Different ionization was allowed within a pH range of 7.0 ± 2.0. All LpxL models were prepared with the Schrödinger protein preparation wizard and minimized using the OPLS4 force field, before generating a grid centered around the residues in the hydrophobic tunnel. The minimized and prepared acyl chains were docked with the Glide program with standard precision (SP) mode, resulting in 10 poses per prepared acyl chain and protein model. The resulting poses were ranked based on the Glide emodel score and the five best scored complexes of each prepared acyl chain and protein model were analyzed visually. The complex with the best pose was further minimized with OPLS4 force field, using water as solvent, method PRCG and maximum iterations 2500. PyMOL was used to prepare pictures of the final complex models and ESPript (https://espript.ibcp.fr) [109] to prepare the alignment picture.

### Cell culture

Immortalised BMDM (iBMDM) cells (BEI Resources, NIAID, NIH: Macrophage Cell Line Derived from Wild Type Mice, NR[9456) were grown in Dulbecco’s modified Eagle’s medium (DMEM; Gibco 41965) supplemented with 10% heat[inactivated foetal calf serum (FCS), 100 U/ml penicillin and 0.1 mg/ml streptomycin (Gibco) at 37°C in a humidified 5% CO_2_ incubator.

Carcinoma human alveolar basal epithelial cells (A549, ATCC CCL-185), and human carcinoma of Henrietta Lacks (HeLa, ATCC CCL-2), were maintained in RPMI 1640 (Gibco) tissue culture medium supplemented with 1% v/v HEPES, 10% v/v heat inactivated fetal bovine serum (FBS) and 1% v/v antibiotics (penicillin and streptomycin) at 37 °C, in a humidified 5% CO2 incubator. Cells were routinely tested for *Mycoplasma* contamination.

### Cell surface expression of TLR4 by flow cytometry

The cell surface expression of TLR4 was determined as previously described [13]. Overnight cultures of *E. coli* MG1655, grown at 37°C in LB, and *Yersinia* strains. grown in LB at 21°C, in an orbital shaker (180 rpm) were diluted 1:10 in LB medium and grown at 21°C or 37°C, *Yersinia* strains, or at 37°C, *E. coli*, for 3.5 h. Bacteria were collected by centrifugation (2500×g, 20 min, 24°C) and resuspended to an OD_600_ of 1.0 in PBS (approximately 5 x 10^8^ CFU/ml). 0.4×10^6^ iBMDMs were seeded in complete DMEM 24-hours prior to lifting with PBS/EDTA and resuspension in complete DMEM. The following day, 0.5×10^6^ iBMDMs were lifted and resuspended in 5 ml tubes in 2 ml of complete DMEM and infected with live bacteria at a multiplicity of infection of 50 bacteria per cell. Following a 20-minute incubation at 37°C, 5% CO2, tubes were placed on ice, centrifuged at 400 x g (4°C), washed with 2 ml of cold PBS, resuspended in 2 ml of ice-cold PBS and stained with antibodies to TLR4 (Biolegend, 117606, 1:200 dilution) for 20 min at 4°C. Cells were then washed with 2 ml ice-cold PBS and centrifuged 400 xg for 5 min, before resuspension in 200 μl PBS. TLR4 monomer surface expression was measured using PE laser channel on the FACs Canto II and analysed using FlowJo version 10.7.2. Data analysis was performed using Prism.

### LPS purification

LPSs from *Yersinia* strains were extracted and purified as previously described using the hot phenol-water method [110]. Bacteria were cultured overnight in 5 ml LB at 21°C. Then, the entire culture was used to inoculate a 2 l flask, containing 1 l of LB broth, which was incubated for 24 hours in agitation at 37 °C. On the next day, an aliquot of the cultures was plated onto LB agar to check for any contamination. Then, cells were collected by centrifugation (6,000 x *g*, for 20 minutes at 4°C). The bacterial pellet was washed twice in PBS, weight and resuspended in deionized water (250 ml of water per 100 g of pellet). Then, the same volume of 65°C pre-warmed liquid phenol was added, and stirred at 65°C in a water bath for 20 minutes. The mixture was then incubated overnight at 4°C following a rapid cool down on ice. Later, samples were centrifuged using a high-speed centrifuge and special glass tubes (30 ml COREX tubes). The aqueous upper phases were combined and transferred into a new 250 ml glass flask. 5 volumes of methanol were added per volume of sample (5:1), and sodium acetate-saturated methanol to a final concentration of 1 % (v/v). Sample was incubated at -20°C overnight. On the next day, samples were centrifuged (6,000 x g, 15 minutes, 4°C). The pellets were resuspended in 20 ml of water, transferred to a dialysis tube and incubated in a tank containing deionized water during 2 days at 4°C, changing the water after 24 h. The content of the dialysis tubes was transferred to a glass flask and lyophilized.

Crude LPS preparations were dispersed (10 mg/ml) in 0.8% NaCl-0.05% NaN_3_-0.1 M Tris-HCl (pH 7), digested with nucleases (50 μg/ml; 1 h 37 °C during with agitation) and proteinase K (50 μg/ml; 3 hours at 55°C and overnight at room temperature). The proteinase K treatment was repeated twice.

LPSs were sedimented by ultracentrifugation (6 h, 100,000 × *g*), and freeze-dried. Protein contamination was tested using a BCA assay following the manufacturer’s recommendations. (Thermofisher). LPSs were repurified using phenol reextraction in the presence of deoxycholate to eliminate lipoprotein contaminants [111].

### Macrophage culture and TNFα ELISA

iBMDMs were seeded 24 h before infection in 24-well plate at a density of 0.5 x 10^8^ cells per well in complete medium. Before infection, cells were washed with PBS and suspended in complete medium without antibiotics. Bacteria were grown in 5 ml LB at 37°C or 21°C, harvested at exponential phase (2,500[×[*g*, 20 min), and adjusted to an OD_600_ of 1.0 in PBS. Infections were performed using a multiplicity of infection of 25 bacteria per cell. Infections were performed in the same medium used to maintain the cell line without antibiotics and incubated at 37°C in a humidified 5% CO_2_ incubator. Cells were centrifuged (200 x g, 5 min) to synchronize infection. After 30 min, cells were washed with PBS and incubated in 100 μg/ml of Gm in complete medium without antibiotics for 3 hours. Supernatants from infected cells were collected and spun down at 12,000[×[*g* for 5 min to remove any debris and stored at -80°C. TNFα in the supernatants was determined using a Murine TNF-α standard 3,3′,5,5′-tetramethylbenzidine (TMB) enzyme-linked immunosorbent assay (ELISA) development kit (PeproTech, catalog number 900-T54).

To challenged iBMDMs with LPSs, cells were seeded the day before treatment in 96-well plates at a density of 0.5×10^5^ cells per well in complete medium. The day of the experiment, cells were washed with PBS and resuspended in complete medium. Cels were challenged with 10 ng/ml of repurified LPSs in complete medium with antibiotics. After 1 h, supernatants were collected, spun down, and stored at -80°C. TNFα was determined using an ELISA (PeproTech, catalog number 900-T54).

### Antimicrobial peptides susceptibility

To assay *Yersinia* strains for resistance to polymyxin B and magainin II, we used a modified version of the sensitivity assay described by Llobet *et al* [49]. Briefly, each strain was grown in LB until an OD_600_ of 0.3, washed once in PBS and diluted in liquid testing media (1% v/v Tryptone soy broth, 10% v/v 100 mM phosphate buffer [pH 6.5], 2% v/v 5 M NaCl) to an approximate concentration of 4 × 10^4^ CFU/ml. Twenty[five microlitres of each diluted strain was then mixed with 5 μl of polymyxin B (10 μg/ml) or magainin II (10 μg/ml) and incubated at 37°C for 1 h. Fifteen microlitres of the suspension was thereafter spread onto LB agar and incubated overnight at 21°C or 37°C. Colony counts were determined and results were expressed as percentages of the colony count of bacteria not exposed to antibacterial agents. All experiments were done with duplicate samples on at least three independent occasions.

### Reporter strains

Two reporter plasmids were constructed to assess the expressions of *inv*, and *ylpA*. The promoter regions of *inv*, and *ylpA* were amplified by PCR using Phusion High-Fidelity DNA Polymerase (NEB), YeO8 genomic DNA as template and the primers shown in Table S3, which contain an EcoRI site at their 5’. The PCR products were gel-purified, digested with EcoRI and cloned into EcoRI-SmaI-digested, Antarctic Phosphatase-treated pGPL01Tp suicide vector [23] and then transformed into *E. coli* GT115 cells to obtain pGPLTp*inv*, and pGPLTp*ylpA*. pGPL01Tp contains a promoterless firefly luciferase *lucFF* gene and an R6K origin of replication and can be mobilized by conjugation [23]. Correct insertion of the amplicons was verified by restriction digestions with EcoRI and SmaI. Vectors were introduced into *E. coli* β2163, and then mobilised into *Y. enterocolitca* strains by conjugation. Cultures were then serially diluted and checked for Tmp resistance by plating on LB Tmp agar at 21°C. Correct insertion of the vectors into the chromosome was confirmed by PCR using the relevant lucFF_check and promoter sequence primers (Table S3).

### Luciferase assays

Overnight cultures of the reporter strain in LB at 21°C, were refreshed 1:10 and grown for 3 h in LB at 21°C in agitation (180 rpm). The cells were then pelleted, washed once in sterile PB buffer (1M disodium hydrogen orthophosphate, 1.5 mM potassium dihydrogen orthophosphate, pH 7.0) and adjusted to an OD_600_ of 1.0. One hundred microlitres of each suspension were added to an equal volume of luciferase assay reagent (1 mM d[luciferin [Synchem] in 100 mM sodium citrate buffer pH 5.0), vortexed for 5 s and then immediately measured for luminescence (expressed as relative light units [RLU]) using a GloMax 20/20 Luminometer (Promega). All strains were tested in quadruplicate from three independent cultures.

### Motility assay

Phenotypic assays for swimming motility were initiated by stabbing 2 µl of an overnight culture at the centre of agar plates containing 0.3% agar and 1% tryptone [52]. Plates were analyzed after 24 h of incubation at 22°C and the diameters of the halos migrated by the strain from the inoculation point were compared. Experiments were run in triplicate in three independent occasions.

### Adhesion and invasion of epithelial cells

Overnight cultures of bacteria in LB at 21oC, were refreshed 1:10 in 5-ml LB and grown at 21°C for 3 h. Bacteria were collected by centrifugation (3,220 x *g*, 20 min at room temperature) and resuspended in PBS to an OD_600_ of 1.0. HeLa cells were seeded in 24-well plates at a density of 5[×[10^4^ in complete RPMI medium containing antibiotics. 2 days after seeding, cells were starved for 16 h before infection using RPMI 1640 medium (Gibco 21875) supplemented only with 10[mM HEPES. Before infection cells were washed once with PBS and infected with a multiplicity of infection of 25 to 1. To determine the adhesion of *Yersinia* to cells, after 30 min infection, monolayers were washed seven times with PBS, and cells were lysed in 300 μl 0.5% saponin in PBS. The lysates were serilally diluted in PBS and plated onto LB agar plates incubated at 21°C. To determine the invasion of cells, after a 30 min infection, monolayers were washed twice with PBS and then incubated for an additional 90 min in medium containing Gm (100 µg/ml) to kill extracellular bacteria. This treatment was long enough to kill all extracellular bacteria. After this period, cells were washed three times with PBS and lysed for 5 min in 300 μl 0.5% saponin in PBS and bacteria were plated. Adhesion and invasion are expressed as CFUs per monolayer. Experiments were carried out in triplicate on three independent occasions.

### Analysis of Yops secretion

Overnight cultures of *Y. enterocolitica* strains were diluted 1∶50 into 25 ml of TSB supplemented with 20 mM MgCl_2_ and 20 mM sodium oxalate in a 100-ml flask. Cultures were incubated with aeration at 21°C for 2.5 h, and then transferred at 37°C for 3 h. The optical density at 540 nm of the culture was measured and the bacterial cells were collected by centrifugation at 1500×*g* for 30 min. Ammonium sulphate (final concentration 47.5% w/v) was used to precipitate proteins from 20 ml of the supernatant. After overnight incubation at 4°C, proteins were collected by centrifugation (3000×*g*, 30 min, 4°C) and washed twice with 1.5 ml of water. Dried protein pellets were resuspended in 50 to 80 µl of sample buffer and normalized according to the cell count. Samples were analyzed by sodium dodecyl sulfate-polyacrylamide gel electrophoresis (SDS-PAGE) using 12% polyacrylamide gels and proteins visualized by Coomassie brilliant blue staining.

### Actin disruption by *Yersinia* infection

A549 cells were seeded on 13 mm circular coverslips in 24-well tissue culture plates to 70% confluence. Cells were serum starved 16 h before infection. Overnight cultures of *Y. enterocolitica* strains grown at 21°C were diluted 1∶10 into 5 ml of LB and grown with aeration at 21°C for 1.5 h and then 1 h at 37°C. Bacteria were pelleted, washed once with PBS and resuspended to an OD_600_[= of 1 in PBS. Cells were infected with this suspension at a multiplicity of infection of 25∶1. After 1 h incubation, the coverslips were washed three times with PBS and then cells fixed with 3.7% PFA in PBS pH 7.4 for 20 min at room temperature. PFA fixed cells were incubated with PBS containing 0.1% saponin, 10% horse serum, Hoechst 33342 (1∶2500), and Orange-Red-phalloidin (1∶100) (Invitrogen) for 30 min in a wet dark chamber. Finally, coverslips were washed twice in 0.1% saponin in PBS, once in PBS and once in H_2_O, mounted on Prolong gold antifade oil (Molecular Probes P36930). Coverslips were visualised on the Leica SP8 confocal microscope. Experiments were carried out in duplicate in three independent occasions.

### Analysis YadA production

Bacteria were grown overnight in 2 ml RPMI 1640 medium lacking phenol red at 37°C without shaking. The OD_540_ of the culture was measured and CFUs were determined by plating serial dilutions. Bacteria from 1-ml aliquot were recovered by centrifugation (16 000×*g*, 10 min, 4°C) and resuspended in 200 µl of SDS-sample buffer. Samples were incubated for 4 h at 37°C and kept frozen at −20°C. Samples were analyzed by SDS-PAGE using 12% polyacrylamide gels and proteins visualized by Coomassie brilliant blue staining. Samples were normalized according to the cell count and they were not boiled before loading the gel.

### Binding assay to collagen-coated slides

Overnight cultures of *Y. enterocolitica* strains grown at 37°C were diluted 1∶10 into 5 ml of LB and grown with aeration at 37°C for 2.5 h. bacteria were pelleted, washed once with PBS and resuspended to an OD_540_ of 0.3 in PBS. 13 mm circular coverslips in 24-well tissue culture plates were coated overnight at 4°C with 10 µg/ml human collagen type IV (Sigma) in PBS (final volume 100 µl). Coverslips were washed three times with TBS and later they were blocked for 1 h at 4°C with 2% BSA in TBS. Finally, coverslips were washed three times and were incubated at 37°C with 100 µl of the bacterial suspension. After 1 h incubation, the coverslips were washed three times with PBS and then bacteria fixed with 3.7% paraformaldehyde (PFA) in PBS pH 7.4 for 20 min at room temperature. PFA fixed cells were incubated with PBS containing 0.1% saponin, 10% horse serum and Hoechst 33342 (1∶25000) for 30 min in a wet dark chamber. Finally, coverslips were washed twice in 0.1% saponin in PBS, once in PBS and once in H_2_O, mounted on Aqua Poly/Mount (Polysciences). Fluorescence images were captured using the ×100 objective lens on a Leica DM5500 microscope equipped with the appropriate filter sets. Acquired images were analysed using the LAS imaging software (Leica). Bacteria were counted in images from three randomly selected fields of view.

### In vivo experiments

8- to 12-week-old Balb/c mice (Charles River) of both sexes were infected by oral gavage with ∼1 × 10^9^ *Y. enterocolitica* strains in 100 μl PBS. 72 h post infection, mice were euthanized using a Schedule 1 method according to UK Home Office approved protocols. Peyer’s patches (3-5 per mouse) were immersed in 1 ml sterile PBS on ice and processed for quantitative bacterial culture immediately. Samples were homogenised with a Precellys Evolution tissue homogenizer (Bertin Instruments), using 1.4 mm ceramic (zirconium oxide) beads at 4,500 rpm for 7 cycles of 10 s, with a 10-s pause between each cycle. Homogenates were serially diluted in sterile PBS and plated onto *Yersinia* selective agar medium plates (Oxoid), and the colonies were enumerated after incubation at 21°C. Data are expressed as CFUs per gr of tissue.

### Statistical analysis

Statistical analyses were performed using one-way analysis of variance (ANOVA) with the Dunnett’s multiple comparisons test, the one-tailed t test, or, when the requirements were not met, the Mann-Whitney U test. Normality and equal variance assumptions were tested with the Kolmogorov-Smirnov test and the Brown-Forsythe test, respectively. Results are given as means ± SD. A *P* value of <0.05 was considered statistically significant. All analyses were performed using GraphPad Prism for Windows (version 10.4.1) software.

## ACKNOWLEDGEMENTS

We are grateful members of Bengoechea lab for helpful discussions. We also thank the bioinformatics (J.V. Lehtonen), and structural biology (FINStruct) infrastructure support from Biocenter Finland and CSC IT Center for Science for computational infrastructure support at the Structural Bioinformatics Laboratory, ACbo Akademi University. The funders had no role in study design, data collection and analysis, decision to publish, or preparation of the manuscript. This work has been funded by Medical Research Council (MRC, MR/V032496/1) and Biotechnology and Biological Sciences Research Council (BBSRC, BB/P006078/1) grants to J.AB. P.J.B. and J.K.M. would like to acknowledge BII core funds. The computational work for this article was partially performed on resources of the National Supercomputing Centre, Singapore (https://www.nscc.sg). This work was funded by the Sigrid Juselius Foundation to T.A.S. and K.M.D.

## SUPPLEMENTARY FIGURES

**Figure S1. Structural characterization of *Yersiniae* lipid A enzymes.**

Negative ion MALDI□TOF mass spectrometry spectra of lipid A purified from (A) *E. coli* BN2 complemented with *lpxM* from *Y. bercovieri,* (B) *E. coli* BN2 complemented with *lpxM* from *Y. intermedia,* (C) YeO8 *lpxP* mutant complemented with *lpxP* from *Y. bercovieri* grown at 21°C, (D) YeO8 *lpxP* mutant complemented with *lpxP* from *Y. intermedia* grown at 21°C, (E) *E. coli* BN1 *lpxL* mutant complemented with *lpxL* from YeO8, (F) *E. coli* BN1 *lpxL* mutant complemented with *lpxL* from *Y. bercovieri*, (G) *E. coli* BN1 *lpxL* mutant complemented with *lpxL* from *Y. molaretii*. Data represent the mass□to□charge (*m/z*) ratios of each lipid A species detected and are representative of three extractions.

**Figure S2. Alignment of *Yersiniae* LpxL homologs.**

The residues are numbered according to YeO8 LpxL. Conserved residues are highlighted in green and residues contributing to the hydrophobic tunnel are marked with black triangles under the alignment. The leucine (pathogenic *Yersinia*) to phenylalanine (non-pathogenic *Yersinia*) mutation, which enables binding of C12 and C14 to LpxL in non-pathogenic strains, is highlighted in pink. The residues that affect the conformation of this phenylalanine in *E. coli* and makes *E. coli* LpxL selective for only C12 are highlighted in yellow. There are four conserved motifs shared with the GPAT family (highlighted in cyan); HX4D/E (motif 1), a conserved arginine (motif 2), a conserved negatively charged residue (motif 3), and a conserved proline (motif 4).

**Figure S3. Modelling the lipid binding site in LpxL.**

(A) *E. coli* LpxL with C12 (orange sticks) bound in the hydrophobic tunnel. The suggested hydrocarbon ruler, F143 (pink sticks), is located in the bottom of the tunnel and interacts with the carbon tail of C12 to form a stable complex. F143 is unlikely to swing away to make space for C14 due to the more polar environment created by Q142. (B) Comparison of C12 (orange sticks) and C14 (light pink sticks) in the hydrophobic tunnel of *E. coli* LpxL. The suggested hydrocarbon ruler, F143 (pink sticks), restricts the tunnel space, causing C14 to stick out into the main groove and a productive complex for catalysis is unlikely to be formed. (C) YeO8 LpxL with C14 (light pink sticks) bound in the hydrophobic tunnel. L146 (pink sticks) is the suggested hydrocarbon ruler and its smaller side chain creates space for C14 to bind, while making it unable to form proper interactions with the shorter C12 for a stable complex. (D) YeO8 and *Y. aldovae* LpxL with C12 (orange sticks) and C14 (light pink sticks) bound in the hydrophobic tunnel. The amino acid numbering is according to Y. aldovae and the only difference in the tunnel is the hydrocarbon ruler (pink sticks); L146 for YeO8 LpxL and F142 for Y*. aldovae* LpxL. F142 in *Y. aldovae* LpxL can probably swing away to create more space in the tunnel and provide a stable interaction site for C14, since the surrounding amino acids L112, I141 and L253 are small hydrophobic residues, which can interact with both F142 and the carbon tail of C14.

**Figure S4. Structural characterization of the lipid A produced by YeO8NPL.**

Negative ion MALDI□TOF mass spectrometry spectra of lipid A purified from (A) YeO8NPL grown at 21°C, (B) YeO8NPL grown at 37°C. Data represent the mass□to□charge (*m/z*) ratios of each lipid A species detected and are representative of three extractions.

**Figure S5. Effect of the lipid A PAMP on *Yersinia* virulence.**

(A) Actin disruption by *Yersinia* infection. A549 cells were non-infected (Control), and infected with YeO8, YeO8 *yopE* mutant (Δ*yopE*, strain YeO8-Δ*yopE*), YeO8NPL, and YeO8 *lpxR* mutant (Δ*lpxR*, strain YeO8-Δ*lpxR*). After fixing and permeabilization of cells actin was stained with Orange-red-phalloidin (1∶100) and cells were analyzed by fluorescence microscopy. Result is representative of three independent experiments. (B) Quantification of the percentage of A549 cells showing actin disruption. Cells were non-infected (Control), and infected with YeO8, YeO8 *yopE* mutant (Δ*yopE*, strain YeO8-Δ*yopE*), YeO8NPL, and YeO8 *lpxR* mutant (Δ*lpxR*, strain YeO8- Δ*lpxR*). The number of cells analysed are indicated on top of each of the bars. (C) Assessment of binding of YeO8, YeO8NPL, and YeO8 cured of the pYV virulence plasmids to collagen-coated coverslips. (D) Quantification of the number of bacteria bound to collagen-coated coverslips.

Data are presented as mean ± SD (n□=□3). In all panels, \*\*\*\**P*□≤□0.0001; ns, *P*□>□0.05 for the comparisons against the results of YeO8 using One□way ANOVA with Dunnett’s multiple comparisons test.

