## Supplementary figures and images for "Interrogating the genus *Yersinia* to define the rules of lipopolysaccharide lipid A structure associated with pathogenicity"

### Fig S1

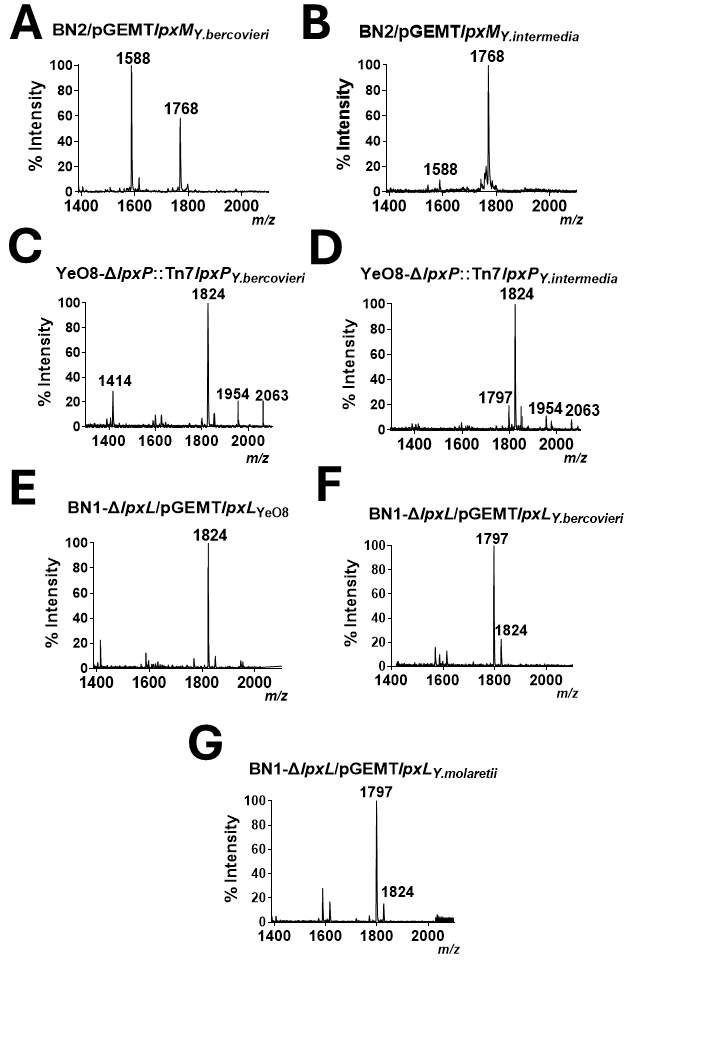

### Fig S2

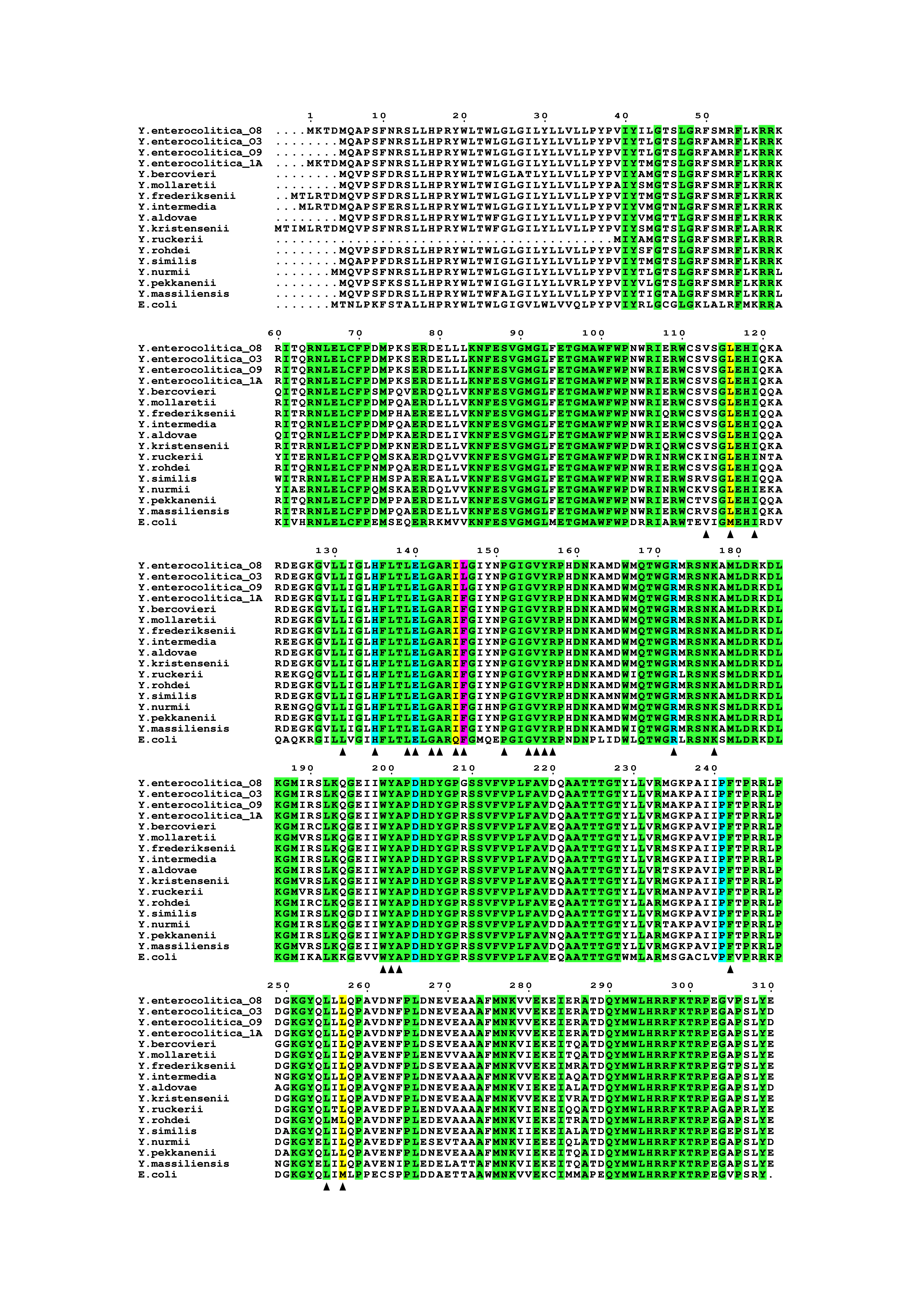

### Fig S3

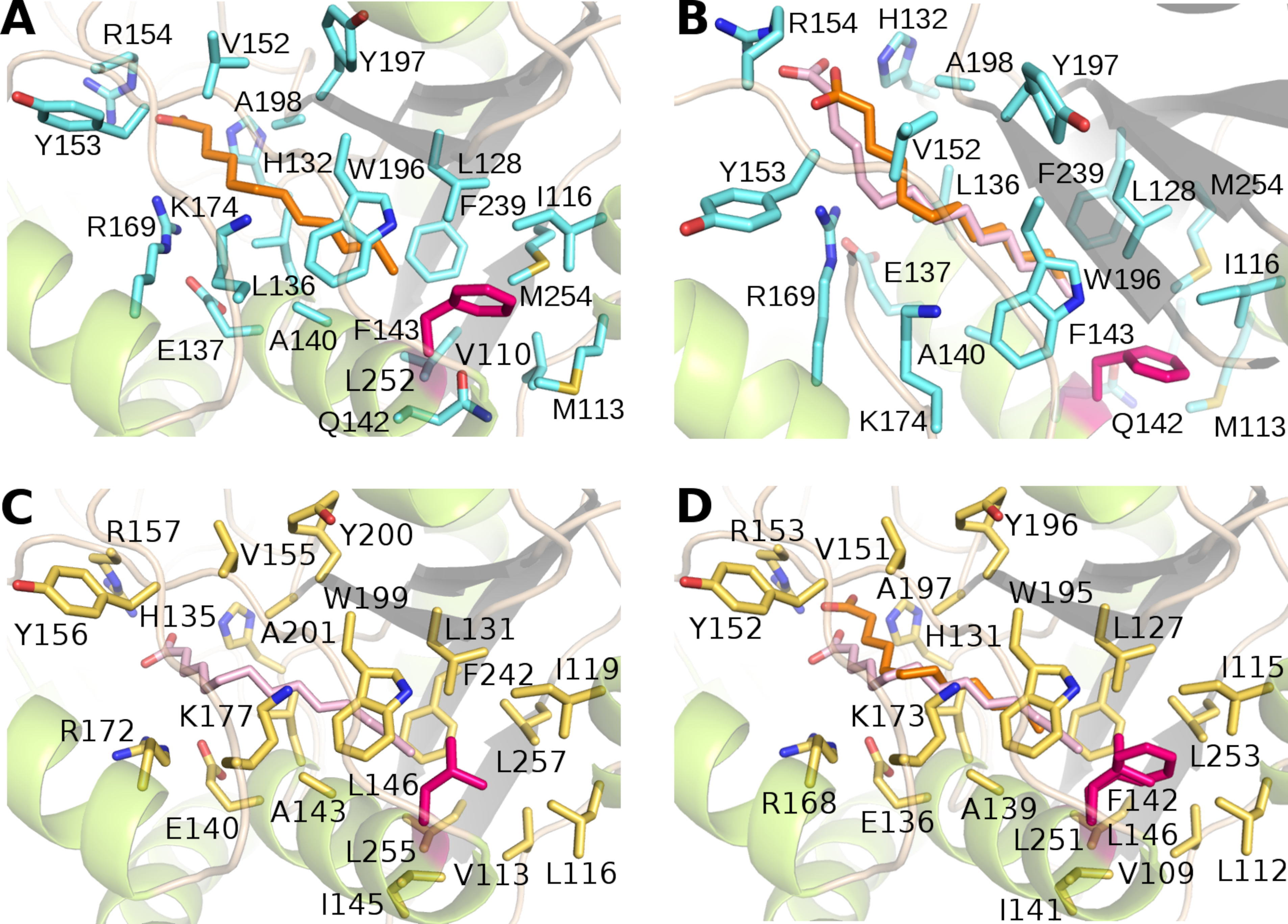

### Fig S4

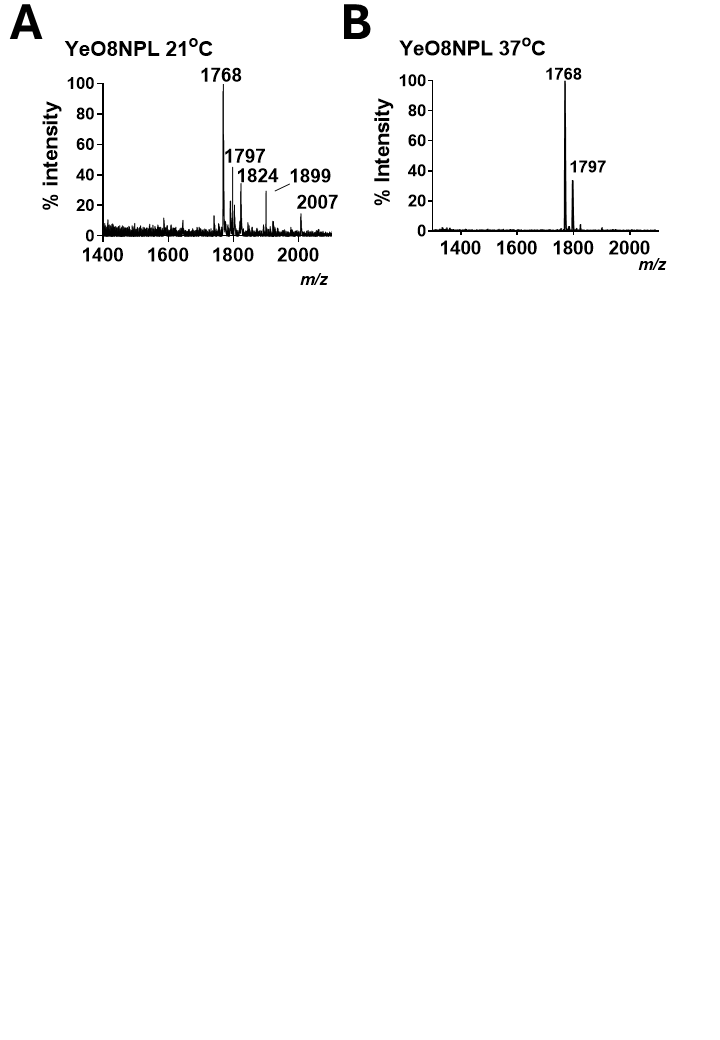

### Fig S5

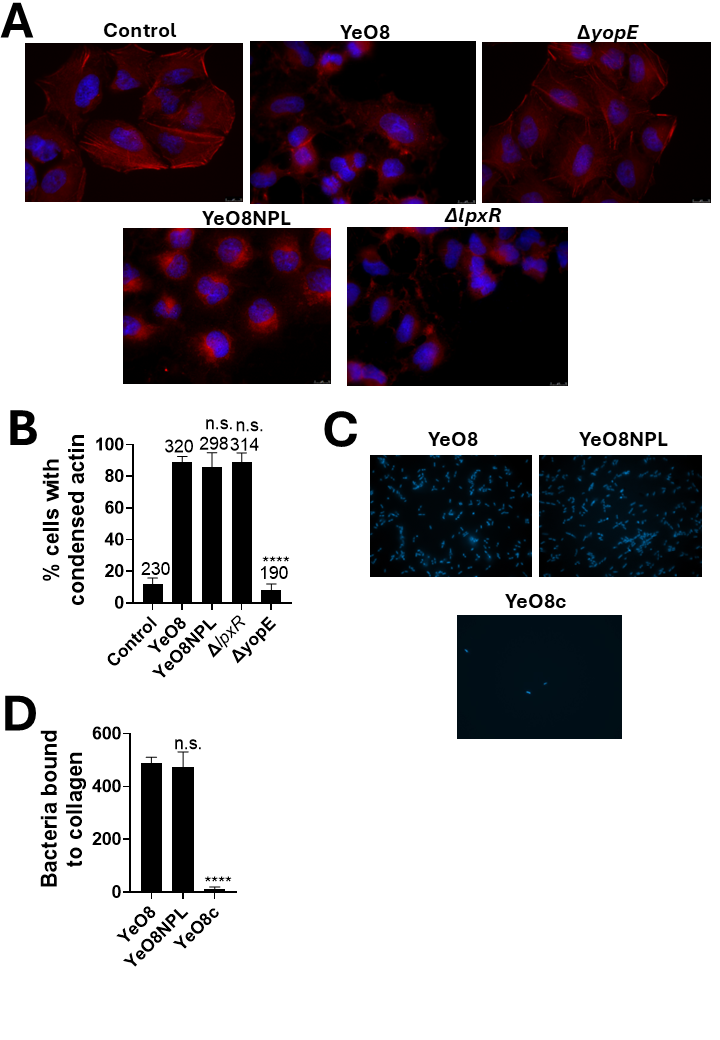
